# Cell-Type-Specific Transcriptomic Signatures in Prefrontal Cortex Reveal lncRNA-mediated Gene Silencing and Enhancer Activities Contributing to the Pathophysiology of Major Depressive Disorder

**DOI:** 10.1101/2025.10.16.682876

**Authors:** Aleena Francis, Yogesh Dwivedi

## Abstract

Long non-coding RNAs (lncRNAs) have emerged as pivotal epigenetic regulators that influence chromatin structure, gene expression, and various cellular processes. We conducted a comprehensive transcriptomic analysis to explore the regulatory roles of lncRNAs in gene expression across major cell types. Based on the cell type abundance as predicted by CIBERSORTx, the gene expression deconvolution of lncRNAs and mRNAs obtained from nonpsychiatric control (n=41) and MDD (n=59) subjects was carried out using bMIND. LncRNA regulatory activity and functional enrichment analysis were assessed in each cell type. Interactions between lncRNAs and transcription factors and RNA-binding proteins, as well as their associations with DNA and RNA, were obtained from the RNA Interactome Database. The protein-protein interactions of lncRNA-regulated mRNAs and the associations between genes and psychiatric disorders were sourced from STRING network PsyGeNET databases, respectively. We observed a prevalence of inhibitory neurons in the control group and a higher proportion of excitatory neurons within the MDD group. A significant dysregulation of lncRNAs and mRNAs, characterized by distinct gene-silencing activity of lncRNAs, was observed, potentially facilitated through chromatin modifications in excitatory and inhibitory neurons, as well as enhancer activity of specific lncRNAs in both neuronal and glial cell types. Functional analyses of the silenced neuronal genes highlighted the involvement of cell adhesion molecules and stress-related signaling in MDD, indicating disrupted neuron-glia interactions. The enhancer activity of lncRNAs indicated the positive regulation of genes involved in Hippo signaling and neurodegeneration. This work uniquely highlights the cell-type-specific roles of lncRNAs and their contributions to MDD pathophysiology.

**HIGHLIGHTS:**

- Comprehensive cell-type-specific transcriptomic analysis of dorsolateral prefrontal cortex in MDD subjects.
- Distinct gene-silencing activity of lncRNAs in excitatory and inhibitory neurons.
- Enhancer activity of specific lncRNAs in both neuronal and glial cell types.
- Critical roles of lncRNAs and associated cellular functions in the pathophysiology of MDD.

## INTRODUCTION

Major Depressive Disorder (MDD) is a pervasive and debilitating condition that affects millions of individuals. The World Health Organization (WHO) has identified MDD as a leading contributor to the global burden of disease, highlighting the urgent need to comprehend its underlying mechanisms and develop more effective therapeutic interventions [1].

Several studies show that stressful experiences can trigger epigenetic modifications, leading to maladaptive responses and increased vulnerability to MDD [2]. Non-coding RNAs, particularly long non-coding RNAs (lncRNAs), have emerged as significant epigenetic modifiers, influencing chromatin structure, gene expression, and various cellular processes [3,4]. LncRNAs are over 200 nucleotides long transcripts that interact with DNA, RNA, and proteins, enabling a wide range of regulatory activities [5]. These versatile molecules can act as scaffolds for protein complexes, guide chromatin-modifying enzymes to specific genomic loci, and influence mRNA splicing and stability. Their involvement in chromatin remodeling, a fundamental process in gene regulation, highlights their importance in cellular differentiation, development, and disease [6]. Specific lncRNAs have been found to be differentially expressed in the brains of MDD subjects, potentially influencing key processes such as neurogenesis, synaptic plasticity, and stress response [7–9]. Moreover, lncRNAs have been implicated in the regulation of genes involved in neurotransmitter signaling, neurotrophic factor expression, and immune function, all of which are relevant to the neurobiology of depression [10]. Studies have revealed distinct sets of lncRNAs and mRNAs associated with depression-susceptible, anxiety-susceptible, and stress-resilient phenotypes in the hippocampus of rats exposed to chronic mild stress [11]. Moreover, our microarray-based transcriptome studies in rat models have elucidated the critical roles of lncRNAs in both susceptibility to and resilience against the development of stress-induced depression, as well as their involvement in the neurobiological response to antidepressant therapies [12]. An extension of this research further underscores the regulatory interactions between mRNA and lncRNAs, highlighting the intricate molecular networks that may inform therapeutic strategies for mood disorders [13]. Additionally, differential lncRNA expression analysis using RNA sequencing of the rostral anterior cingulate cortex, combined with WGCNA-based co-expression analysis, has identified modules of highly co-expressed distally located protein-coding genes associated with lncRNAs in relation to depression and suicide [14].

While the role of lncRNAs in depression has garnered increasing attention, cell type-specific studies are poised to revolutionize our understanding of their functions in this disorder [15]. LncRNAs exhibit notable region- and cell-type-specific patterns, suggesting that their functions may vary across different cell populations in the brain. In the context of MDD, this specificity is crucial because different brain cell types, such as neurons, oligodendrocytes, astrocytes, and microglia, play distinct roles in the pathophysiology of the disorder. For example, lncRNA FEDORA is predominantly expressed in oligodendrocytes and neurons and is upregulated in the prefrontal cortex of depressed females [15]. Furthermore, a single-cell transcriptome analysis revealed that lncRNAs can be highly expressed in individual cells of the developing neocortex and are cell-type-specific [16]. Thus, this level of resolution is essential for understanding how lncRNAs influence cell-autonomous functions, intercellular communication, and the overall neural circuitry implicated in depression [15].

In the present study, we deconvolved the bulk lncRNA and mRNA expression data from dorsolateral prefrontal cortex (dlPFC) using a reference gene expression profile derived from snRNASeq. We found significant differences in cell type proportions between the control and MDD cohorts. Our differential expression analysis revealed dysregulated lncRNAs and mRNAs in MDD. Furthermore, correlation analyses indicated that upregulated lncRNAs play roles in both gene silencing and enhancer activities across major cell types. Altogether, the study uniquely highlights cell-type-specific roles of lncRNAs in MDD pathophysiology, advancing understanding of molecular mechanisms and offering novel targets for therapeutic intervention.

## MATERIALS & METHODS

### Postmortem human brain samples

The Institutional Review Board of the University of Alabama at Birmingham approved the study. Samples from dlPFC were sourced from the Maryland Brain Collection at the Maryland Psychiatric Research Center in Baltimore, MD. The cohort included 100 participants, consisting of 59 individuals diagnosed with MDD and 41 non-psychiatric control subjects. Brain pH was assessed using established protocols [17]. Comprehensive demographic and clinical information about the subjects is presented in **Table S1**. Age, PMI, brain pH, and RNA integrity number (RIN) were not significantly different between the control and MDD groups. There were 26 males and 15 females in the control group and 35 males and 24 females in the MDD group. Detailed inclusion-exclusion criteria and brain sample collection are described in the **supplementary section**.

### Microarray analysis

Total RNA was extracted from the dlPFC utilizing TRIzol (Invitrogen) according to the manufacturer’s instructions provided earlier [18] and detailed in the **supplementary section**. Expression of lncRNAs and mRNAs was assessed using a custom-designed microarray platform, which included probes for 40,000 annotated human lncRNAs and 30,000 mRNAs. Total RNA (500 ng) was labeled using the Agilent Quick Amp Labeling Kit (Agilent Technologies) and hybridized to the microarray slides according to the manufacturer’s instructions. Slides were scanned using an Agilent G2565CA Microarray Scanner System, and raw intensity data were extracted using Agilent Feature Extraction software.

### Data preprocessing, cleansing, and normalization

The raw microarray data were preprocessed and normalized using the Limma package in R, as previously outlined [19]. Initially, background correction was applied, followed by quantile normalization to standardize expression values across samples. Probes with low signal intensities, specifically those below the 20th percentile, were excluded from the analysis. The expression data for every sample were log2-transformed for subsequent study. In total, 19,757 mRNAs and 35,003 lncRNAs were detected.

### snRNASeq data analysis

The reference gene expression profile was constructed de novo using snRNASeq data from GSE144136 processed through the Seurat package (v5.2.1) [20]. The snRNASeq data was separated into control and MDD conditions, and the cells were filtered for nFeature_RNA>400 & nFeature_RNA<5000 & nCount_RNA>1000. The cell-type clustering was performed using principal component analysis (PCA) and t-Distributed Stochastic Neighbor Embedding (tSNE) for visualization. Cell types were assigned based on prior annotations available in the GEO dataset. The heatmap of the marker genes for each cell type was plotted using pheatmap R package [21].

### Cell Type deconvolution

The cell type deconvolution was carried out using CIBERSORTx (http://cibersortx.stanford.edu) platform. The snRNA-Seq data with annotations were uploaded to the create signature matrix module of CIBERSORTx to generate a new signature matrix for both control and MDD groups using default settings, while ‘Min.Expression’ was set to 0.5, and the option to “Disable quantile normalization” was left unchecked. Subsequently, cell fractions were estimated separately for both groups by submitting the normalized and log-transformed bulk transcriptome data of lncRNA and mRNA to the impute cell fractions module and employing S-mode batch correction with 100 permutations for significance assessment. The association between predicted cell type proportions and demographic variables, including age, PMI, and pH, was assessed using Pearson correlation analysis. Furthermore, differences in cell type proportions associated with categorical variables, such as sex and cause of death, were evaluated utilizing the Wilcoxon rank-sum test.

### Expression deconvolution and differential analysis

Based on the cell type abundance as predicted by CIBERSORTx, the gene expression deconvolution of lncRNAs and mRNAs was carried out using the bMIND R package [22]. The Wilcoxon test for differential expression analysis between MDD and control was performed for both lncRNAs and mRNAs, utilizing the deconvoluted expression dataset obtained on a sample-wise basis. The p-value adjustment was performed using the Benjamini-Hochberg (BH) method [23]. An absolute log2 fold change of ≥ 0.21 and an adjusted p-value threshold of 0.05 were applied to identify significantly differentially regulated genes across the various cell types.

### Analysis of lncRNA regulatory activity

The Pearson correlation between mRNA and lncRNA expression in each cell type was calculated using the cor.test() function in R. Significant correlations were identified based on a p<0.05 and an R² value 0.4. For negative regulation, we investigated the correlations between upregulated lncRNAs and downregulated mRNAs having an R² threshold <-0.4. In contrast, for enhancer activity, we analyzed the correlations between upregulated lncRNAs and upregulated mRNAs, focusing on R² values >0.4. The proximity of lncRNA and mRNA pairs was assessed if both transcripts were situated on the same chromosome and were separated by a distance less than 100kb.

### Functional enrichment analysis and tissue-wise expression profiling

The functional enrichment analysis of KEGG pathways and Gene Ontologies (GO), comprising biological process, molecular function, and cellular component, was performed using the clusterProfiler package [24]. The results were visualized using the enrichplot::barplot() function. The expression profile of lncRNAs and their regulated genes was obtained from the GTEx portal (https://www.gtexportal.org/home/) for the selected brain tissues, including amygdala, anterior cingulate cortex, caudate, cerebellar hemisphere, cerebellum, cortex, frontal cortex, hippocampus, hypothalamus, nucleus accumbens, putamen, spinal cord, and substantia nigra.

### lncRNA interaction and localization analysis

Interactions between lncRNAs and various proteins, including transcription factors and RNA-binding proteins, as well as their associations with DNA and RNA, were sourced from the RNA Interactome Database (RNAInter) (http://www.rnainter.org/). The interactions of the top ten lncRNAs, determined by the highest correlation values and exhibiting at least one type of interacting partner, were visually represented in a network format using Cytoscape (v3.10.2). Only the top two interactions with the highest score from each category were included. Additionally, information on the subcellular localization of lncRNAs was collected from the RNALocate (http://www.rnalocate.org/) database.

### Interaction network analysis

The protein-protein interactions of mRNAs regulated by the lncRNAs were extracted from the STRING network database (v12), having a minimum confidence score of 0.4. These interactions were visualized using Cytoscape (v3.10.2). Additionally, the Cytohubba plugin was utilized to identify the top 20 hub genes within each network, ranked based on the Maximal Clique Centrality (MCC) method [25].

### PsyGeNET analysis

The associations between genes and psychiatric disorders were sourced from the PsyGeNET database (http://www.psygenet.org/). We explored mRNAs that are either upregulated or downregulated by the lncRNAs to examine their links to psychiatric conditions such as depression and schizophrenia. All entries from the database were taken into consideration. A score of 1 signifies a 100% positive association, whereas 0 indicates no association with the disorder.

## RESULTS

### Single-nuclei RNASeq analysis

The snRNA sequencing analysis of dlPFC from both control and MDD individuals presents a comprehensive reference gene expression profile for these distinct conditions. The investigation revealed the presence of six primary cell types in both control and MDD samples: excitatory neurons, inhibitory neurons, oligodendrocytes, oligodendrocyte precursor cells (OPCs), astrocytes, and micro/macroglial cells. Notably, excitatory neurons were identified as the predominant cell type, while micro/macroglial cells constituted the least abundant population within both control and MDD cohorts (**Fig. 1A-B**). The expression profile of cell type-specific marker genes provides a robust validation of cell type assignment and underscores the specificity inherent to this classification. Notably, the expression of marker genes such as *SLC17A7, SATB2,* and *NEUROD6* was exclusively associated with excitatory neurons, whereas *GAD1, GAD2*, and *PNOC* were distinctly expressed in inhibitory neurons (**Fig. 1C-D**).

**Fig. 1:**
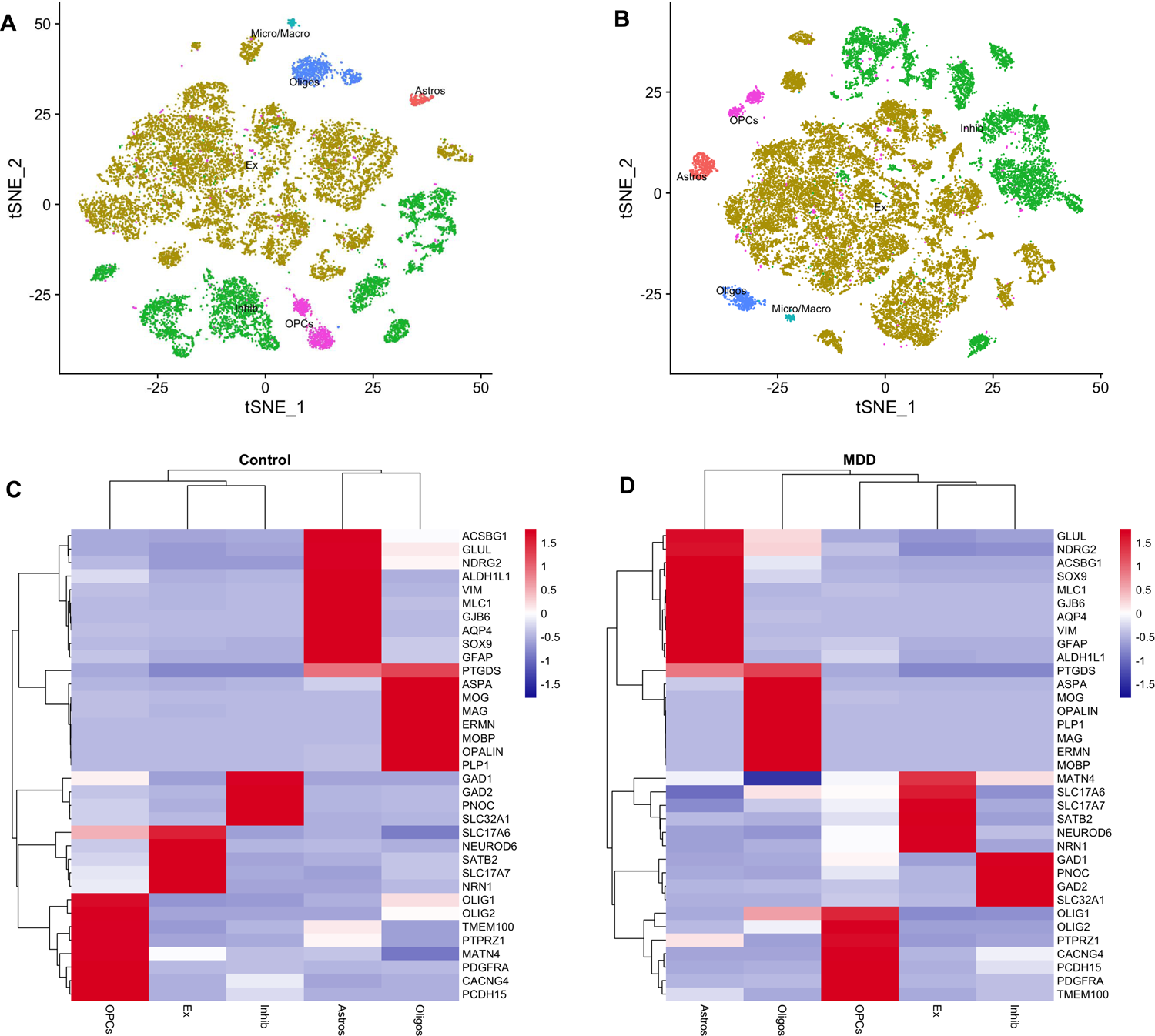
Single-nuclei transcriptomic analysis of brain prefrontal cortex cell populations in Control and MDD subjects. **A-B** t-SNE visualization of single-nuclei transcriptomic data, displaying major brain cell types in **A** Control and **B** MDD samples. Different cell types are color-coded, including oligodendrocytes (Oligo), astrocytes (Astros), microglia/macrophages (Micro/Macro), inhibitory neurons (Inhib), excitatory neurons (Ex), and oligodendrocyte precursor cells (OPCs). **C-D** Heatmaps of canonical cell-type specific marker gene expression across brain cell types in the scRNASeq data of **C** Control and **D** MDD conditions. Gene expression is Z-score normalized, with red indicating high and blue indicating low expression values. Rows represent genes, and columns represent different cell types, as indicated at the bottom of the heatmaps. Clustering dendrograms illustrate the similarity between gene expression patterns across cell types.

### snRNASeq reference gene matrix using CIBERSORTx

We generated a custom signature expression profile for the control and MDD groups using the reference gene expression profile from the snRNASeq data. The heatmap produced by CIBERSORTx for the signature matrix illustrates the specificity of gene expression for individual cell types in both groups (**Fig. S1A-B**).

### Prediction of cell type proportion in bulk transcriptome

The abundance of individual cell types in the bulk transcriptome was determined using CIBERSORTx with the signature gene expression matrix. A total of 54,760 microarray probes corresponding to mRNAs and lncRNAs were used for the cell type deconvolution (**Table S2-3**). Notably, control samples exhibited a significant enrichment of inhibitory neurons, while the MDD cohort demonstrated a marked enrichment of excitatory neurons. Additionally, the astrocyte and oligodendrocyte fractions were substantially reduced in the MDD samples, suggesting potential alterations in glial cell populations associated with this condition (**Fig. S1C-D**). These findings align with prior studies implicating glial dysfunction in the pathophysiology of MDD and highlight potential disruptions in neuron-glia interactions [26,27]. The correlation between cell-type marker gene expression and cell-type abundance further corroborates the accuracy of the deconvolution method (**Fig. S2**). Age and sex distributions were significantly associated with the proportion of oligodendrocytes in MDD. All other demographic features showed no significant association with the proportions of other cell types (**Fig. S3**).

### Expression deconvolution using bMIND

Based on the cell type proportion, expression deconvolution was carried out using bMIND, as it provides a consolidated expression profile for individual cell types, as well as sample-wise expression for differential expression analysis. The cell-type marker genes *SATB2, GAD2, GFAP, PCDH15,* and *MOBP* for the excitatory neurons, inhibitory neurons, astrocytes, OPCs, and oligodendrocytes, respectively, exhibited the strongest correlation between the expression data derived from snRNASeq data and deconvoluted expression profile (**Fig. S4**).

### Differential Expression of lncRNAs and mRNAs Across Cell Types

The differential expression analysis of lncRNAs and mRNAs was conducted across five major cell types. The analysis revealed variability in the number of differentially expressed lncRNAs and mRNAs among the distinct cell types. The highest number of dysregulated lncRNAs was found in inhibitory neurons (2,300), followed by astrocytes (1,194), while excitatory neurons had the lowest (198). Upregulated lncRNAs were most prevalent in inhibitory neurons (748) and astrocytes (575). (**Fig. 2A, Table S4**). Among the 198 lncRNAs that were differentially regulated in excitatory neurons, 134 were unique to this cell type. Approximately 92% of the lncRNAs were specific to inhibitory neurons, while 199 were specific to OPCs, 147 to oligodendrocytes, and 1,017 to astrocytes.

**Fig. 2:**
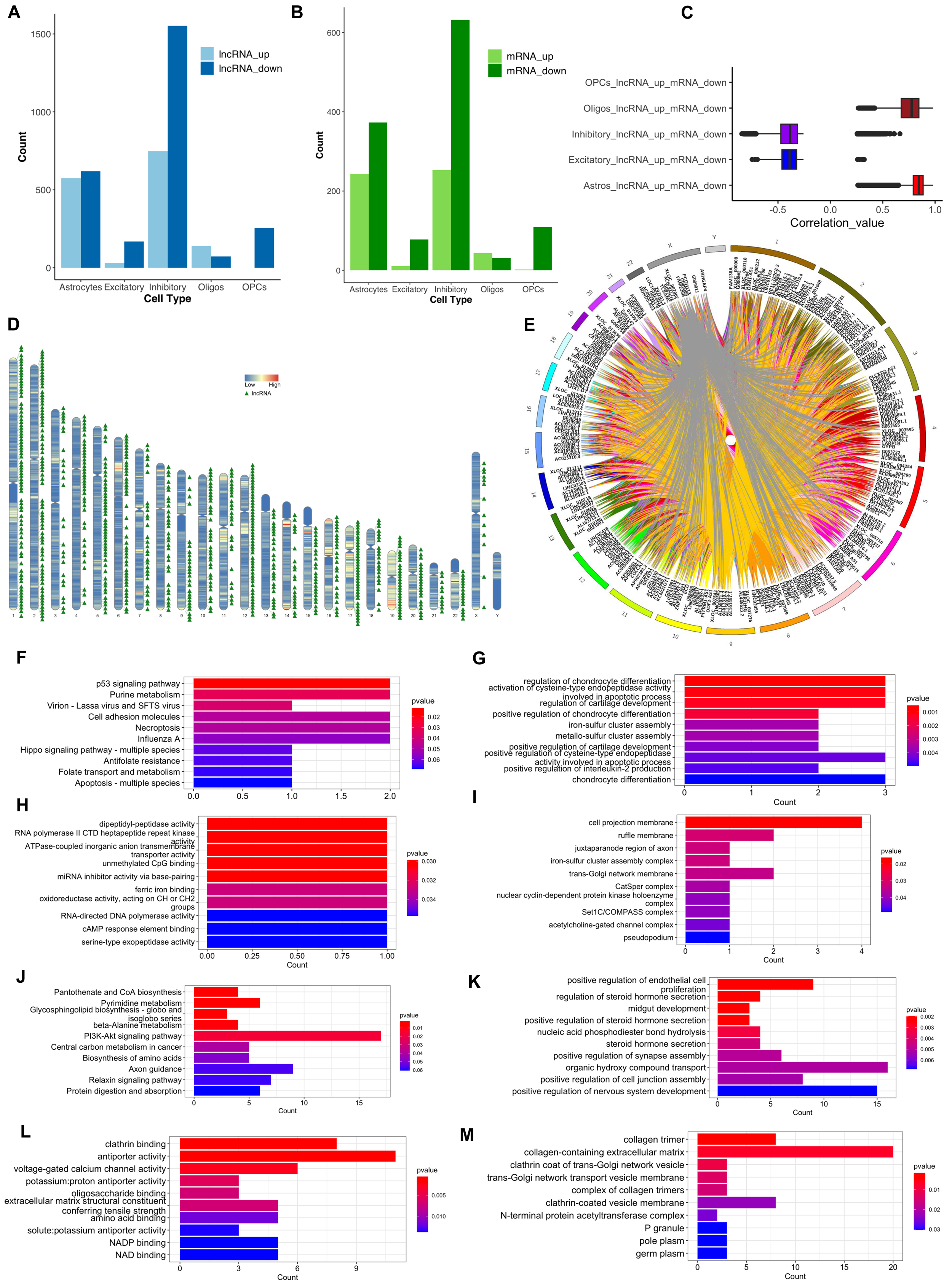
The analysis of differential expression, the inhibitory activity of lncRNAs on mRNAs, and the regulatory effects of lncRNAs in inhibitory neurons. **A** A bar graph representing the number of up- and downregulated lncRNAs across four cell types: Astrocytes, Excitatory neurons, Inhibitory neurons, and OPCs. **B** A bar plot representing the number of differentially expressed mRNAs (upregulated and downregulated) in different cell types. **C** The Pearson correlation values between the upregulated lncRNAs and downregulated mRNAs across various cell types are illustrated as a box plot. This analysis explores the relationships between lncRNA and mRNA expression, highlighting a potential negative regulatory role of upregulated lncRNAs on downregulated mRNAs, as indicated by negative correlation values. Notably, such negative correlations were observed only in excitatory and inhibitory neurons. **D** Chromosomal distribution of the upregulated lncRNAs with negative regulatory effect on mRNAs identified in inhibitory neurons. The karyotype is depicted with blue and red indicating low and high gene density, respectively. **E** The circos plot depicts the genomic distribution of negatively correlated upregulated lncRNAs and downregulated mRNAs in inhibitory neurons. The outer ring represents the chromosomes, while the inner connections indicate pairs of correlated lncRNAs and mRNAs. The functional analysis of the negatively correlated up-lncRNAs and down-mRNAs in inhibitory neurons. **F** KEGG pathway enrichment of the lncRNAs. **G** Biological Process GO term enrichment of the lncRNAs. **H** Molecular Function GO term enrichment of the lncRNAs. **I** Cellular Component GO term enrichment of the lncRNAs. **J** KEGG pathway enrichment of the mRNAs. **K** Biological Process GO term enrichment of the mRNAs. **L** Molecular Function GO term enrichment of the mRNAs. **M** Cellular Component GO term enrichment of the mRNAs.

Similarly, the total differentially expressed mRNAs range from 75 in Oligos to 885 in inhibitory neurons, with inhibitory neurons having the highest number of upregulated (253) and downregulated (632) mRNAs. Notably, astrocytes showed a substantial number of differentially expressed genes, totaling 616 mRNAs (243 up-regulated and 373 down-regulated). The least number of differentially regulated mRNAs was observed in oligodendrocytes (75), followed by excitatory neurons (89) and OPCs (111) (**Fig. 2B, Table S5**).

### Cell-type-specific gene regulation by lncRNAs in dlPFC

#### a. Gene silencing activity

The Pearson correlation analysis between the cell-type-specific expression of mRNAs that are down-regulated and lncRNAs that are up-regulated in the MDD group reveals the negative regulatory effect of lncRNAs in silencing transcription activity by possible heterochromatin formation (**Fig. 2C**). Negative regulation of lncRNA on gene expression in MDD was observed in both inhibitory and excitatory neurons, but not in other cell types.

##### i. Inhibitory neurons

Interestingly, the highest number of correlating pairs was observed in inhibitory neurons, which included 739 lncRNAs and 627 mRNAs (**Table S6**). The distribution patterns of lncRNAs in the genome reveal that Chr2 harbors the highest abundance of lncRNAs. Notably, these lncRNAs are predominantly concentrated within the q arm of the chromosome, indicating a potential correlation between lncRNA distribution and genomic organization (**Fig. 2D**). The nuclear localization of 148 lncRNAs was identified, underscoring their significance in establishing interactions with the DNA and modifying the chromatin environment (**Table S7**). Additionally, all of the lncRNAs and mRNAs associated with silencing activity were identified as primarily functioning through trans-regulatory mechanisms based on the genomic location except for eight lncRNA-mRNA pairs displaying evidence of proximal regulation through a distance <100kb (*CATG00000018224.1-RPGRIP1*:Chr14, *TERC-LRRIQ4*:Chr3, *G042637-HS1BP3*:Chr2, *AC096920.1-C3orf18*:Chr3, *CATG00000088168.1-CLPS*:Chr6, *PRSS16-POM121L2*:Chr6, *AC022211.1-GGA3*:Chr17, and *CATG00000055826.1-MAP3K7CL*:Chr21) (**Fig. 2E**).

###### Functional analysis of lncRNAs

The upregulated lncRNAs from inhibitory neurons displayed a significant enrichment in several key biological pathways and molecular functions. Notably, there was a marked enrichment of cell adhesion molecules and pathways associated with the p53 signaling cascade (**Fig. 2F**). In terms of biological processes, critical regulatory functions, including the activation of cysteine-type endopeptidase activity pertinent to the apoptotic process, and the positive regulation of interleukin-2 production, were highlighted (**Fig. 2G**). Additionally, molecular function terms such as miRNA inhibitor activity, characterized by base pairing, unmethylated CpG binding, and RNA-directed DNA polymerase activity, were identified (**Fig. 2H**). Furthermore, the cellular component analysis revealed an enrichment in cell projection membranes, underscoring the multifaceted roles of these lncRNAs in neurobiological contexts (**Fig. 2I**).

###### Functional analysis of mRNAs

The mRNAs potentially silenced by the lncRNAs exhibited an enrichment of pathways implicated in the PI3K-Akt signaling cascade, pyrimidine metabolism, and amino acid biosynthesis (**Fig. 2J**). Significant biological processes identified included organic hydroxy compound transport, positive regulation of synapse assembly, and positive regulation of nervous system development (**Fig. 2K**), while prominent molecular functions highlighted were antiporter activity, clathrin binding, and voltage-gated calcium channel activity (**Fig. 2L**). Enrichment was also observed in cellular components such as the collagen-containing extracellular matrix and clathrin-coated vesicle membranes, further emphasizing the intricate nature of lncRNAs in cellular regulation (**Fig. 2M)**.

###### Tissue-wise expression analysis

The gene expression profiles of lncRNAs and their associated mRNAs were analyzed using the GTEx database. The lncRNAs LINC00667, SNX17, and LYRM4 displayed consistently elevated expression across all examined brain tissues (**Fig. 3A**). The mRNAs that are regulated by these lncRNAs, including LENG8, TBCB, PGK1, FAM131A, and DNAJC6, exhibited higher expression across all brain areas while, the genes including COL6A5, MYOC, DLX3, and SLC9A2 were characterized by notably lower expression levels throughout the examined regions (**Fig. 3B**).

**Fig. 3:**
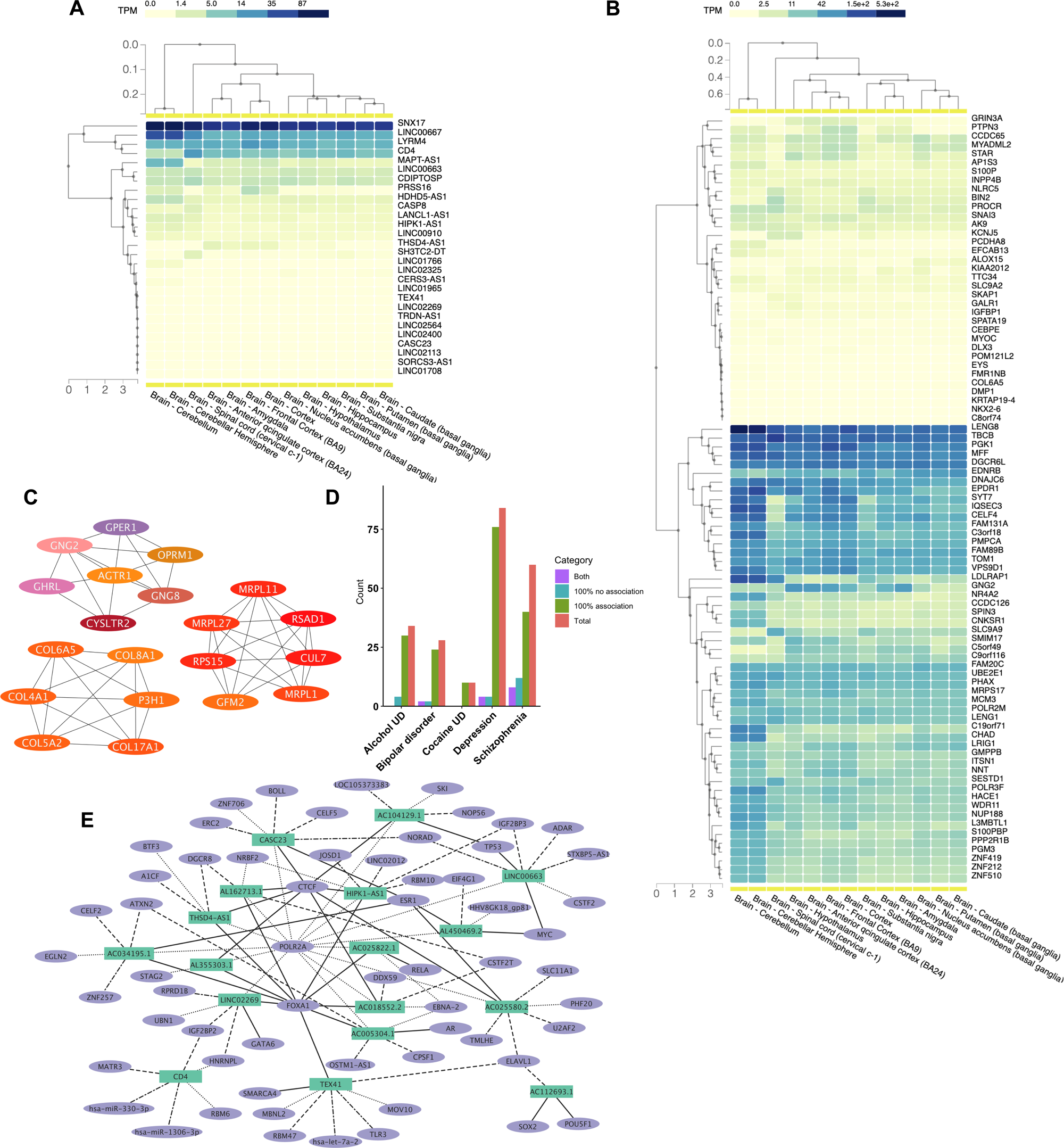
Functional characterization of the dysregulated correlating pairs of lncRNA and mRNAs in inhibitory neurons exhibiting inhibitory activity. **A** A heatmap delineating the expression levels (TPM) of selected lncRNAs originating from inhibitory neurons across various brain regions, as obtained from the GTEx database. **B** A heatmap delineating the expression levels (TPM) of selected mRNAs originating from inhibitory neurons across various brain regions, as obtained from the GTEx database. **C** Three major protein-protein interaction networks of the 20 hub genes identified among the negatively regulated mRNAs in inhibitory neurons. **D** The number of genes associated with various neurological disorders from PsyGeNET database, including Alcohol Use Disorder, Bipolar Disorder, Cocaine Use Disorder, Depression, and Schizophrenia, identified among the negatively regulated mRNAs in inhibitory neurons. 100% no association indicates that the genes are not linked to the disorder, while 100% association shows that the genes are positively linked to the disorder. Both means there are reports of both positive and no association. **E** Network representation of known interactions between the top lncRNAs in inhibitory neurons and DNA, RNA, and proteins from the RNAInter database. Only the top two interactions with the highest score from each category are included. In the diagram, interactions with DNA are represented as parallel lines, RNA interactions as dash-dot lines, protein interactions as dots, RNA binding proteins as dashed lines, and transcription factors as solid lines.

###### PPI network analysis

The protein-protein interaction network analysis of the potentially regulated transcripts revealed three distinct networks comprising 20 hub genes. These hub genes included extracellular matrix organization genes such as COL8A1, COL6A5, COL4A1, and COL5A2, as well as translation-related genes, specifically MRPL11, MRPL27, MRPL1, RPS15, and GFM2. Additionally, genes associated with the GPCR ligand binding pathway, including GPER1, GNG2, GHRL, CYSLTR2, AGTR1, GNG8, and OPRM1, were identified among the hub genes (**Fig. 3C**).

###### PsyGeNET analysis

The established relationship between genes and psychiatric disorders enhances our understanding of the implications these genes have in such conditions. A significant number of genes that demonstrate a negative correlation with lncRNAs were found to have a positive association with depressive disorders (76), followed by schizophrenia (40) and alcohol use disorder (30) (**Fig. 3D**).

###### lncRNA interaction analysis

The experimentally validated interactions of lncRNAs with various biomolecules, including RNAs, DNAs, and proteins such as transcription factors (TFs) and RNA-binding proteins (RBPs), offer valuable insights into the underlying mechanisms through which lncRNAs exert their regulatory effects on gene expression. Among the up-regulated lncRNAs, 44 demonstrated interactions with a total of 1,472 unique gene loci. Additionally, 178 lncRNAs interacted with 928 RNA molecules, while 378 exhibited associations with 13,251 proteins. Furthermore, 366 lncRNAs interacted with 276 RBPs, and 345 demonstrated interactions with 758 TFs (**Table S8**). The analysis of interactions among the lncRNAs exhibiting the highest correlation values revealed that the transcription factor FOXA1 engaged with ten specific lncRNAs, including AC005304.1, AC018552.2, AC025822.1, AC034195.1, AL162713.1, AL355303.1, HIPK1-AS1, LINC02269, TEX41, and THSD4-AS1.

Additionally, the CTCF TF was found to interact with the lncRNAs AC018552.2, AC025580.2, AC104129.1, AL162713.1, AL355303.1, CASC23, HIPK1-AS1, and THSD4-AS1. Notably, fourteen lncRNAs exhibited interactions with the polymerase II (POLR2A) protein (**Fig. 3E**).

##### ii. Excitatory neurons

Among the excitatory neurons, a significant negative correlation was observed between 609 pairs comprising 29 upregulated lncRNAs and 69 downregulated mRNAs, elucidating the complex regulatory landscape associated with this cell type (**Table S6**). The analysis of chromosomal distribution revealed that chromosomes 4, 8, 12, 13, 21, 22, X, and Y were devoid of any differential lncRNAs, while Chr 2 exhibited the highest abundance (**Fig. S5A**). The nuclear localization of twelve lncRNAs was validated through the RNALocate database, lending support to their potential involvement in the modulation of chromatin structure (**Table S7**). Additionally, all inhibitory lncRNAs and mRNAs were located distally, either on the same or different chromosomes, suggesting complex gene regulation mechanisms. (**Fig. S5B**).

###### Functional analysis of mRNAs

The functional analysis of mRNAs that are negatively regulated revealed a significant enrichment in pathways associated with focal adhesion, the p53 signaling pathway, and resistance to EGFR tyrosine kinase inhibitors (**Fig. S5C**). Furthermore, biological process terms such as cellular response to extracellular stimuli, regulation of cell cycle phase transitions (**Fig. S5D**), associated molecular functions including enzyme inhibitor activity, rRNA binding, and protein methyltransferase activity (**Fig. S5E**), and pore complex cellular component were found to be enriched among the genes (**Fig. S5F**).

###### Tissue-wise expression analysis

The evaluation of gene expression in different brain tissues indicates their specific expression patterns. The lncRNAs LINC00963, IDI2-AS1, and ATP6V0E2-AS1, were highly expressed across all brain regions examined. Notably, HOTAIRM1 showed expression exclusively in the spinal cord, with no detectable levels in other brain tissues (**Fig. S6A**). The genes that are negatively regulated by these lncRNAs, such as NGRN, IGIPFAM168, ARHOJ, and ZNF207, showed consistent expression across all brain regions studied. In contrast, other genes, including PCDHA5, ZNF852, CDKN2B, and FAS, had inherently lower expression. (**Fig. S6B)**.

###### PPI network analysis

An analysis of protein-protein interactions demonstrated significant networks among negatively regulated genes in excitatory neurons. Key interactions identified include BCL2 with CDKN2B, C5, and FAS; RHOJ with TRIO and USP24; ACO1 with ETFA; and BRIX1 interacting with PINX1 (**Fig. S6C**).

###### PsyGeNET analysis

The association of dysregulated genes with psychiatric disorders revealed a positive connection of eight genes with schizophrenia and alcohol use disorder, followed by four genes linked to bipolar disorder and two to depression (**Fig. S6D**).

###### lncRNA interaction analysis

Analysis of known interactions of the lncRNAs having inhibitory activity with other biomolecules indicates that eleven lncRNAs were observed to interact with 2,186 RNAs, 19 interact with 6,108 proteins, 16 are associated with 233 RBPs, 20 with 604 TFs, and one lncRNA interacts with ten different gene loci (**Table S9**). The HOXA1 TF RNA and protein were found to interact with HOTAIRM1 lncRNA. Eukaryotic translation initiation factors EIF3D and EIF4G1 are interacting with LINC01135 and AC093904.2, respectively. Furthermore, the CTCF TF exhibited interactions with LINC00608, LINC00963, AC093904.2, LINC01135, PLCG1-AS1, AC127496.3, ATP6V0E2-AS1, HOTAIRM1, and LINC00506 lncRNAs, distinct from inhibitory neurons (**Fig. S6E**).

#### b. Enhancer activity

We investigated the positive regulatory effects of lncRNAs on mRNAs by performing correlation analyses between the expression levels of upregulated lncRNAs and mRNAs. Notably, enhancer activity was observed across all examined cell types, except for OPCs (**Fig. 4A**). In excitatory neurons, a total of 20 unique lncRNAs were found to be associated with eight mRNAs. While in inhibitory neurons, we identified 723 lncRNAs linked to 252 mRNAs. Oligodendrocytes exhibited associations between 138 lncRNAs and 44 transcripts, whereas astrocytes demonstrated a positive correlation between 561 lncRNAs and 243 mRNAs (**Table S10**).

**Fig. 4:**
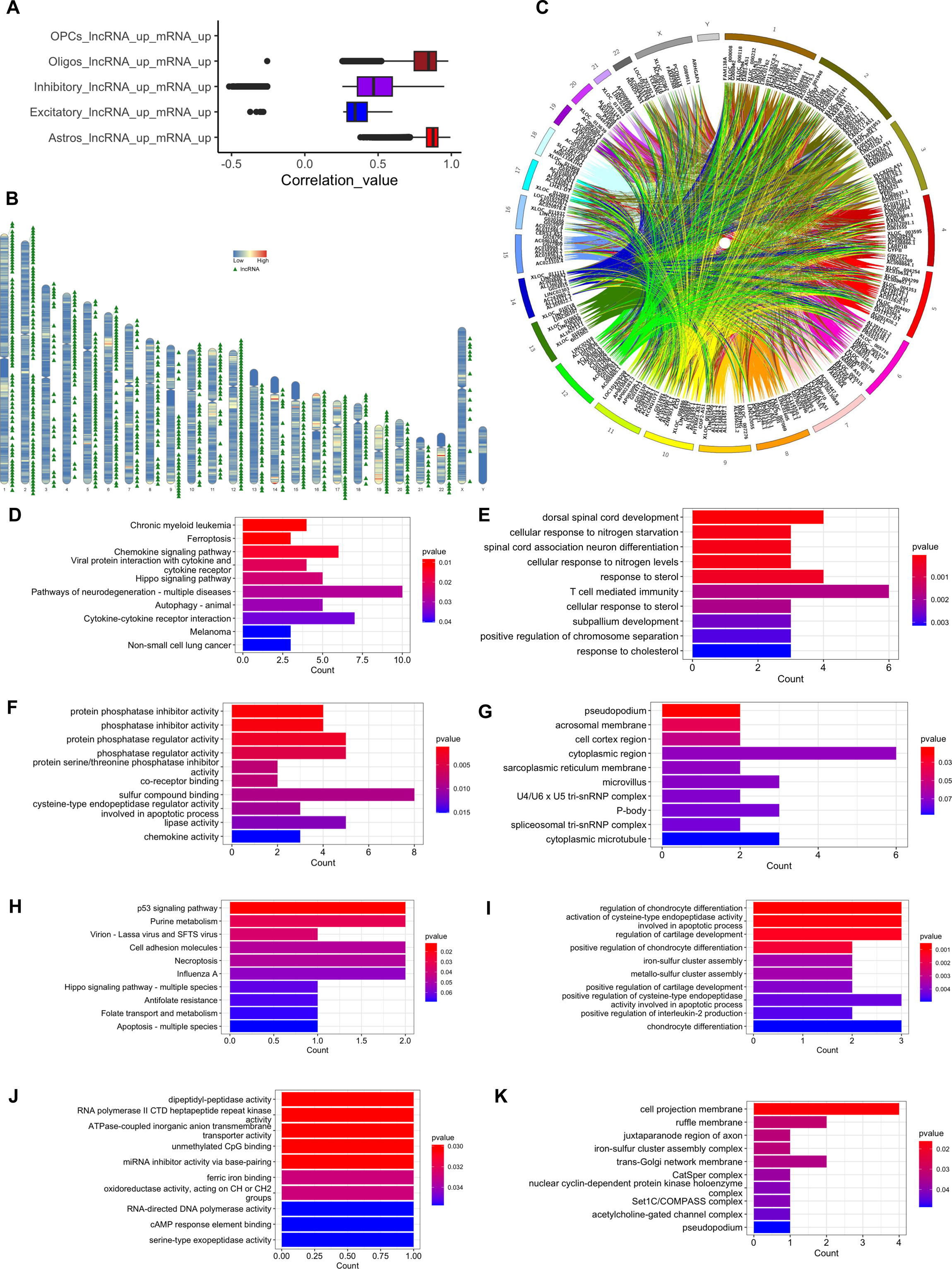
The analysis of the enhancer activity of lncRNAs on mRNAs, and the regulatory effects of lncRNAs in inhibitory neurons. **A** The Pearson correlation values between the dysregulated lncRNAs and mRNAs across various cell types are illustrated using a box plot. This analysis explores the relationships between lncRNA and mRNA expression, highlighting a potential positive regulatory role of upregulated lncRNAs on upregulated mRNAs, as indicated by positive correlation values. **B** Chromosomal distribution of the upregulated lncRNAs with positive regulatory effect on mRNAs identified in inhibitory neurons. The karyotype is depicted with blue and red indicating low and high gene density, respectively. **C** The circos plot depicting the genomic distribution of positively correlated upregulated lncRNAs and upregulated mRNAs in inhibitory neurons. The outer ring represents the chromosomes, while the inner connections indicate pairs of correlated lncRNAs and mRNAs. **D** KEGG pathway enrichment of the positively correlated mRNAs in inhibitory neurons. **E** Biological Process GO term enrichment of the positively correlated mRNAs in inhibitory neurons. **F** Molecular Function GO term enrichment of the positively correlated mRNAs in inhibitory neurons. **G** Cellular Component GO term enrichment of the positively correlated mRNAs in inhibitory neurons. **H** KEGG pathway enrichment of the lncRNAs. **I** Biological Process GO term enrichment of the lncRNAs. **J** Molecular Function GO term enrichment of the lncRNAs. **K** Cellular Component GO term enrichment of the lncRNAs.

##### i. Inhibitory neurons

The unique lncRNAs that are positively associated with gene expression in inhibitory neurons are distributed across all chromosomes, with the highest numbers found on Chr1 (73) and Chr2 (66) (**Fig. 4B**). The lncRNAs AL592463.1, AC073342.1, AC138150.2, AL603832.2, AL139300.2, and AL021707.7 are located near the associated transcripts MUSK, TAS2R39, ACBD4, FAM19A3, TRMT61A, and FAM227A, respectively. All other lncRNA-transcript pairs were either located on different chromosomes or were not within a proximity of 100 kb (**Figure 4C**).

###### Functional analysis of mRNAs

The functional analysis of the upregulated mRNAs associated with enhancer lncRNAs showed an enrichment in the Hippo signaling pathway, neurodegeneration pathway (**Fig. 4D**), and biological processes including T cell-mediated immunity and spinal cord-associated neuron differentiation (**Fig. 4E**). In terms of molecular functions, there was enrichment for sulfur compound binding, protein phosphatase regulator activity, and lipase activity (**Fig. 4F**). Additionally, the genes were enriched for the cytoplasmic region (**Fig. 4G**).

###### Functional analysis of lncRNAs

The functional enrichment analysis of the enhancer lncRNAs reveals significant associations with the p53 signaling pathway, cell adhesion molecules, and necroptosis pathways (**Fig. 4H)**. Key biological processes include the regulation of chondrocyte differentiation and interleukin-2 production (**Fig. 4I**). Molecular functions observed involve miRNA inhibition, binding to cAMP response elements, and interaction with unmethylated CpG motifs (**Fig. 4J**). Notably, the cell projection membrane exhibited the highest enrichment among cellular components (**Fig. 4K**).

###### Tissue-wise expression analysis

A detailed assessment of the expression profiles of the mRNAs revealed that genes such as MLC1, G6PC3, and ZBTB47 exhibited higher expression levels across multiple brain regions. In contrast, WNT3A, TRIM42, ADM2, and LIPN demonstrated markedly lower expression levels. (**Fig. 5A**). Moderate expression levels of DNAJC9-AS1 and consistently lower expression of lncRNAs LINC01940, LINC00358, LINC02420, LINC02564, and LINC02269 were observed in all brain areas (**Fig. 5B**).

**Fig. 5:**
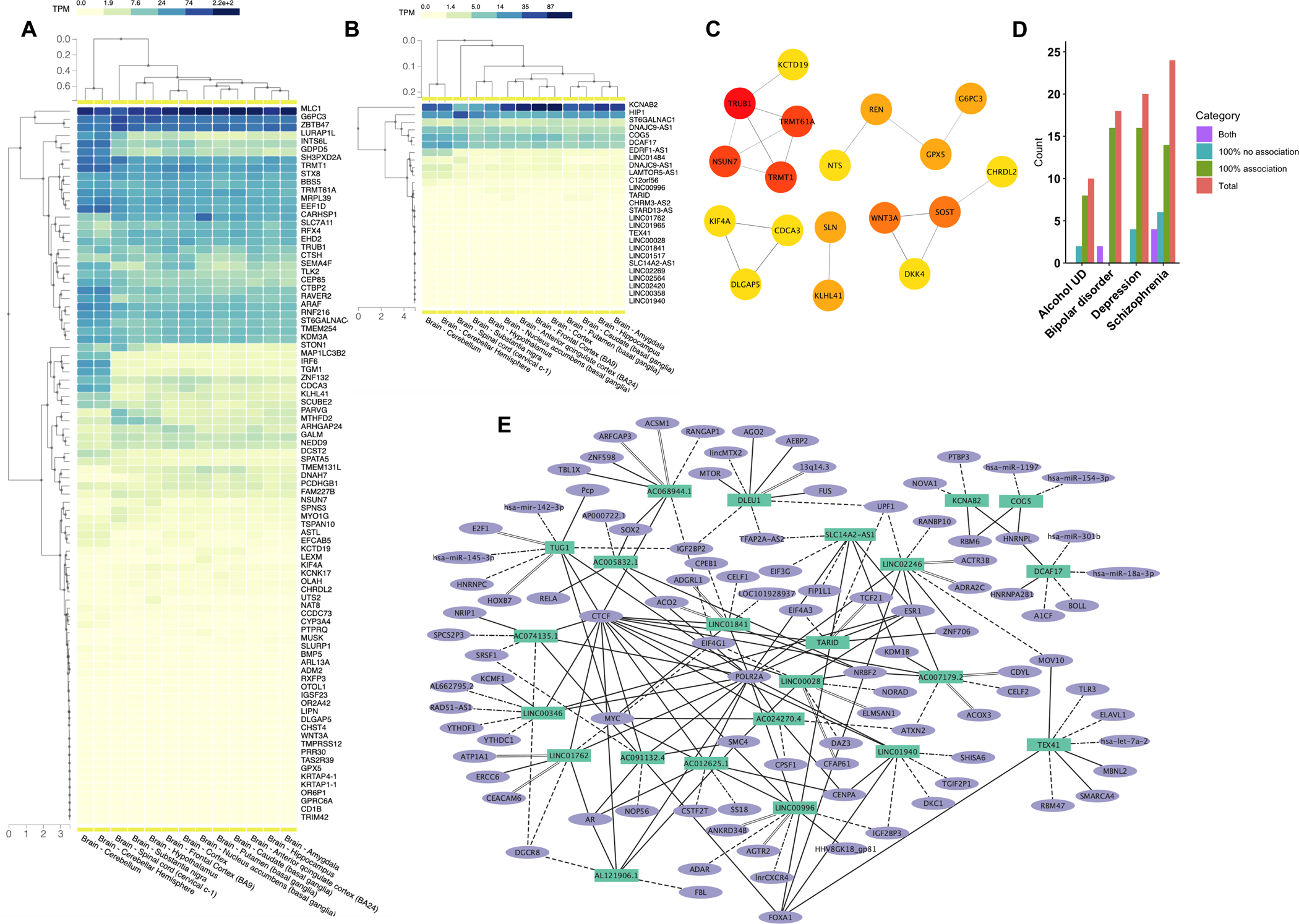
Functional characterization of the dysregulated correlating pairs of lncRNA and mRNAs in inhibitory neurons exhibiting enhancer activity. **A** A heatmap delineating the expression levels (TPM) of selected mRNAs originating from inhibitory neurons across various brain regions, as obtained from the GTEx database. **B** A heatmap delineating the expression levels (TPM) of selected lncRNAs originating from inhibitory neurons across various brain areas, as obtained from the GTEx database. **C** Major protein-protein interaction networks of the 20 hub genes identified among the negatively regulated mRNAs in inhibitory neurons. **D** The number of genes associated with various neurological disorders from PsyGeNET database, including Alcohol Use Disorder, Bipolar Disorder, Depression, and Schizophrenia, identified among the positively regulated mRNAs in inhibitory neurons. 100% no association indicates that the genes are not linked to the disorder, while 100% association shows that the genes are positively linked to the disorder. Both means there are reports of both positive and no association. **E** Network representation of known interactions between the top lncRNAs from inhibitory neurons and DNA, RNA, and proteins from the RNAInter database. Only the top two interactions with the highest score from each category are included. In the diagram, interactions with DNA are represented as parallel lines, RNA interactions as dash-dot lines, protein interactions as dots, RNA binding proteins as dashed lines, and transcription factors as solid lines.

###### PPI network analysis

Among the mRNAs exhibiting enhanced expression mediated by lncRNAs, the STRING clusters associated with the Wnt signaling pathway, specifically WNT3A, DKK4, SOST, and CHRDL2, as well as those involved in tRNA modification and enzyme-directed rRNA pseudouridine synthesis, including TRUB1, NSUN7, TRMT61A, and TRMT1, were identified as hub genes (**Fig. 5C**).

###### PsyGeNET analysis

An analysis utilizing the PsyGeNET database revealed a positive association of 16 genes with depressive disorders and bipolar disorder (**Fig. 5D**).

###### LncRNA interaction analysis

Investigations into the interactions of lncRNAs exhibiting the highest correlation values, SLC14A2-AS1, TARID, and TUG1, have been identified as interacting with the POLR2A protein. Furthermore, lncRNAs AC005832.1, AC007179.2, LINC00028, and LINC01841 are associated with NRBF2. The RNA-binding protein EIF4G1 has demonstrated interactions with AC068944.1, LINC00028, and LINC01762, while RBP IGF2BP2 interacts specifically with DLEU1 and TUG1. Additionally, multiple TFs, including CTCF, FOXA1, and MYC, have shown interactions with a range of lncRNAs, suggesting a complex regulatory network. A cohort of microRNAs, hsa-let-7a-2, hsa-miR-1197, hsa-mir-142-3p, hsa-miR-145-3p, hsa-miR-154-3p, hsa-miR-18a-3p, and hsa-miR-301b has also been implicated in these interactions. Notably, the 13q14.3 chromosomal region is associated with the DLEU1 lncRNA, further underscoring the intricate interplay between lncRNAs and other molecular entities within cellular regulatory mechanisms (**Fig. 5E, Table S11**).

##### ii. Excitatory neurons

The expression correlation between the lncRNA and mRNA revealed that excitatory neurons had the lowest pairs, with only 45 identified. Additionally, the chromosomal distribution analysis indicated that the lncRNAs were restricted to chromosomes 2, 3, 6, 7, 9, 14, 15, 16, 17, 18, 19, and 20 (**Fig. S7A**). Among these, only ATP6V0E2-AS1 and AC124014.1 showed a positive correlation with genes located on the same chromosome but more than 100kb apart; all other pairs were found on different chromosomes (**Fig. S7B**).

###### Functional analysis of mRNAs

The functional enrichment analysis of the upregulated mRNAs revealed several significant pathways and GO terms. Notably, these included the spinocerebellar ataxia pathway and the olfactory transduction pathways (**Fig. S7C**). The analysis highlighted biological processes such as motor neuron migration and the negative regulation of receptor signaling pathways mediated by the JAK-STAT pathway (**Fig. S7D**). Additionally, we identified enriched molecular functions, including serine-type endopeptidase activity and olfactory receptor activity (**Fig. S7E**).

###### Tissue-wise expression analysis

The expression profiling showed that lncRNAs ATP6V0E2-AS1 and LINC00963 were expressed in all brain tissues. (**Fig. S7F)**. The RNF187 gene showed consistently higher expression across the brain tissues, while the DAB1 mRNA showed expression in the cerebellum and cerebellar hemisphere region only (**Fig. S7G**).

###### LncRNA interaction analysis

The analysis of lncRNA interactions elucidated that the POLR2A protein engages with ten distinct lncRNAs, including AC093904.2, ATP6V0E2-AS1, KC6, LINC00506, LINC00608, LINC00661, LINC00963, and PLCG1-AS1. Furthermore, the RBP IGF2BP3 demonstrates interaction with both ATP6V0E2-AS1 and LINC00506. Notably, the TF CTCF is associated with six different lncRNAs: AC093904.2, ATP6V0E2-AS1, LINC00506, LINC00608, LINC00963, and PLCG1-AS1 (**Fig. S7H, Table S12**).

The gene enhancer activity of lncRNAs in oligodendrocytes and astrocytes is discussed further in the **supplementary section**.

## DISCUSSION

Recently, lncRNAs have been recognized as crucial regulators of gene expression, demonstrating significant cell-type specificity within the human brain. In this study, we aim to investigate the unique landscape of lncRNA and gene regulation in PFC cell types using the expression deconvolution technique. This research represents a pioneering effort in the field to study the implications of the lncRNAs on the pathophysiology of MDD. The potential gene-silencing activity of lncRNAs was assessed by analyzing the inverse relationship between upregulated lncRNAs and downregulated mRNAs in MDD. Our study revealed significant negative regulation of lncRNAs in both excitatory and inhibitory neurons, indicating these molecules may play a crucial role in the gene expression dysregulation associated with MDD [28,29].

Functional enrichment analyses highlighted several key biological processes and molecular functions associated with MDD. The enrichment of cell adhesion molecules (CAMs) among dysregulated lncRNAs from inhibitory neurons suggests that disruptions in cell-cell interactions and synaptic connectivity may contribute to the pathophysiology. CAMs are essential for neuronal migration, axon guidance, synapse formation, and synaptic plasticity, all of which are critical for normal brain function [30]. Dysregulation of CAM expression may lead to impaired neuronal connectivity and contribute to the cognitive and emotional symptoms of MDD [31]. The analysis also highlights the regulatory roles of lncRNAs in crucial biological functions such as the activation of cysteine-type endopeptidase activity and the positive regulation of interleukin-2 production. The activation of cysteine-type endopeptidases is pertinent to apoptosis, a mechanism implicated in stress-related neuronal loss [32]. Moreover, the regulation of interleukin-2, a cytokine critical for T-cell proliferation and immune response, suggests that lncRNAs may interface with the immune system in the context of neuroinflammatory processes associated with MDD [9]. Such roles emphasize the interdisciplinary nature of lncRNAs, where immunological and neurobiological angles converge to offer a holistic perspective on MDD pathogenesis. Moreover, pathways associated with the p53 signaling cascade among dysregulated lncRNAs from inhibitory neurons and mRNAs in excitatory neurons underscore the role of cellular stress and apoptosis within neural populations contributing to MDD. The tumor suppressor protein p53 is activated in response to various cellular stresses, including DNA damage, oxidative stress, and inflammation, leading to cell cycle arrest, DNA repair, or apoptosis [33,34]. Activation of the p53 pathway has been implicated in the pathogenesis of MDD, with evidence suggesting that increased apoptosis in key brain regions contributes to the structural and functional abnormalities observed in this disorder [35].

The protein-protein interaction network analysis has revealed a complex landscape of cellular communication and regulatory mechanisms through hub genes, thereby reinforcing the interconnectedness of cellular pathways involved in neurobiology and synaptic regulation. Among the hub genes in inhibitory neurons, several extracellular matrix (ECM) organization genes were highlighted. ECM genes are vital in maintaining the structural integrity and functionality of tissues, playing prominent roles in neuronal architecture [36]. The ECM not only modifies physical tissue properties but also influences neuronal signaling and plasticity, essential for processes such as synapse formation and maintenance [37]. Moreover, the implication of ECM organization in synaptic function suggests that lncRNAs could significantly shape neuronal responses and structural adaptations. In addition to ECM-related genes, the cluster also included translation-related genes, specifically *MRPL11, MRPL27, MRPL1, RPS15,* and *GFM2*. These MRPL proteins are integral components of mitochondrial ribosomes [38], underscoring the importance of mitochondrial function in neuronal health and metabolism. Mitochondrial protein synthesis is crucial for energy production and can influence neuronal growth, differentiation, and synaptic efficacy [39]. Disruption in mitochondrial translation has been linked to neurodegenerative diseases and cellular stress responses, indicating that the regulation of these genes by lncRNAs could have profound effects on neuronal survival and function [40]. Moreover, within the negatively regulated genes from excitatory neurons, *RHOJ*, an important Rho GTPase, connected with processes involving cytoskeletal dynamics and cell motility, interacts with TRIO, a guanine nucleotide exchange factor, showcasing how lncRNA interactions can influence not just existing neuronal pathways but also cellular morphology and migration [41].

The interaction of transcription factor FOXA1 with ten distinct lncRNAs in inhibitory neurons clearly stands out. The engagement of FOXA1, known for its role as a pioneer factor that can open chromatin for transcriptional activation [42], implies that these lncRNAs could potentially influence gene expression patterns linked to development, metabolism, and response to extracellular signals. Also, the involvement of the CTCF TF with multiple lncRNAs points to their shared participation in chromatin structure regulation and gene expression. CTCF, recognized for its role in defining chromatin domains and regulating long-range enhancer-promoter interactions [43], enhances the functional relevance of these lncRNAs in maintaining genomic architecture and influencing transcriptional outcomes during neuronal function and development. In excitatory neurons, both FOXA1 and CTCF have been demonstrated to interact with HOTAIRM1, a significant epigenetic regulator that plays a crucial role in various mechanisms associated with depression. This regulation occurs through the modulation of neurotransmitters, neurotrophic factors, and synaptic functions [44].

The investigation into the positive regulatory effects of lncRNAs on mRNAs presents compelling evidence correlating their expression levels in all neuronal and glial cells. In all cell types, the lncRNAs function in trans at distal genomic locations to regulate the expression of protein-coding genes, with a few notable exceptions in inhibitory neurons and astrocytes. One such pair in inhibitory neurons, the lncRNA AL592463.1, is located near the MUSK gene. The MUSK gene plays a critical role in neuromuscular synapse formation [45]. This genomic arrangement facilitates a targeted and efficient mechanism of gene control, allowing for precise tuning of neuronal activities and synaptic plasticity. Proteins such as POLR2A and the transcription factor CTCF exhibited interactions with numerous lncRNAs with positive regulatory functions in all the cell types analyzed; however, these interactions involved different lncRNAs, suggesting distinct lncRNAs play roles specific to each cell type. The presence of POLR2A binding sites in the promoter regions of various lncRNAs indicates that RNA Polymerase II plays a crucial role in activating the transcription of these lncRNAs [46].

Altogether, our study offers a detailed transcriptomic characterization of MDD by identifying cell-type-specific changes and lncRNA-mediated gene regulation. These are marked by distinct gene-silencing functions of lncRNAs, likely influenced by chromatin modifications in both excitatory and inhibitory neurons, as well as the enhancer roles of select lncRNAs in neuronal and glial cell populations. Through an integrated analysis of gene expression, functional enrichment, protein-protein interactions, and relationships to psychiatric disorders, we have highlighted key molecular pathways and potential therapeutic targets for MDD. Our findings support the importance of cell-type-specific analysis in understanding the complex neurobiology of psychiatric conditions and promote further investigation into the therapeutic potential of lncRNAs.

## ACKNOWLEDGMENTS

YD is the Elesabeth Ridgley Endowed Chair and is supported by National Institute of Mental Health grants R01MH130539, R01MH124248, R01MH118884, R01MH128994, R01MH107183, and R56MH138596.

## AUTHOR CONTRIBUTIONS

YD obtained the funding, conceptualized the study, and supervised the study design and implementation. AF performed the deconvolution analysis and wrote the first draft. YD finalized the manuscript.

## MATERIALS & CORRESPONDENCE

No materials were generated in the study. Correspondence and material requests should be addressed to Yogesh Dwivedi, Ph.D.

## DATA AND CODE AVAILABILITY

All data supporting the findings of this study are available within the article and its supplementary files. Any additional requests for information can be directed to and will be fulfilled by the corresponding author. This paper does not report the original code.

## Supplementary Methods

### Postmortem Human Brain Samples

Tissue samples from control subjects and MDD patients were screened for neuropathology and excluded if they showed signs of Alzheimer’s disease, infarcts, demyelination, atrophy, or a related clinical history. Urine and blood samples were analyzed for toxicology and antidepressant presence. The psychiatric diagnosis was determined by psychological autopsy as described earlier using Diagnostic Evaluation After Death (DEAD) [1] and the Structured Clinical Interview for the DSM-5 (SCID) [2]. The interviews were conducted by a qualified psychiatric social worker. Two psychiatrists separately assessed the write-up from this interview, along with the SCID that was completed, as part of their diagnostic evaluation of the case. Diagnoses were derived from the information gathered during this interview, the medical records related to the case, and reports from the Medical Examiner’s office. The two diagnoses were compared, and any discrepancies were addressed through a consensus conference. Control subjects were confirmed to be free from mental illnesses by using these consensus diagnostic methods.

After removal from the cranium, the brains were cut into six major pieces (four cerebral cortical lobes, basal ganglia-diencephalon, and lower brain stem-cerebellum), rapidly frozen on dry ice, and stored at −70°C until dissection. During dissection, the frontal lobes were sliced into 1-mm to 1.5-mm thick coronal sections at a temperature between 0°C and 10°C. To keep the samples frozen, the dissections were performed on a metal plate over a container filled with dry ice. The prefrontal cortical samples were cut out of the coronal sections by a fine microdissecting (Graefe) knife under a stereomicroscope with low magnification. The dorsomedial prefrontal cortex (Brodmann’s area 9) was taken just dorsal to the frontopolar area, including the most polar portion of the superior and partly the middle frontal gyrus between the superior and intermediate frontal sulci. In the sections of the dissected cortical area, the gray and white matter were separated. The tissues were chopped into smaller pieces and stored at −80°C until use. The tissues were examined histologically. Fixed sections of PFC were screened with hematoxylin and eosin (H&E) staining and an antibody to glial fibrillary acid protein.

### RNA Isolation

Roughly 50 mg of tissue was homogenized in 1 mL of TRIzol reagent with the help of a motorized homogenizer. Following chloroform extraction, the aqueous phase was obtained, and RNA was precipitated using isopropanol. The RNA pellet was washed with 75% ethanol, air-dried, and then resuspended in RNase-free water. The quality and integrity of the RNA were evaluated with an Agilent Bioanalyzer 2100, and samples with an RIN of 7 or higher were chosen for subsequent analyses.

## Supplementary Results

### a. Enhancer activity

#### i. Oligodendrocytes

A total of 5,824 pairs exhibiting a minimum correlation coefficient of 0.4 were identified within the oligodendrocyte cell type, encompassing 138 unique lncRNAs and 44 mRNAs. The chromosomal distribution shows that Chr8 has the highest number of lncRNAs, followed by Chr7 and Chr2. Additionally, 6 lncRNAs are located on chromosome X (**Fig. S8A**). Of the total pairs identified, 288 were located on the same chromosome, while the remainder were distributed across different chromosomes (**Fig. S8B**).

##### Functional analysis of lncRNAs

The enriched lncRNAs highlighted pathways such as glycophingolipid biosynthesis, along with biological processes including maintenance of apical/basal cell polarity. Additionally, pre-mRNA 5’ splice site binding molecular function, and the cellular component associated with the voltage-gated calcium channel complex were identified (**Fig. S8C-F)**.

##### Functional analysis of mRNAs

The genes that are positively regulated by the lncRNAs showed enrichment in the TNF signaling pathway and the calcium signaling pathway, as well as activities linked to guanyl-nucleotide exchange factors, small GTPase-mediated signal transduction, regulation of canonical NF-kappaB signal transduction, and microtubule-related cellular components (**Fig. S8G-J**).

##### Tissue-wise expression analysis

The expression levels of lncRNAs LINC00957 and RABL2A and mRNAs PAQR3, ARHGEF40, and PHKG2 were found to be consistently higher across all examined brain-related tissues (**Fig. S9A-B**).

##### PPI network analysis

Significant protein-protein interactions were identified among the following gene pairs: DUSP19 and PAQR3, TRPA1 and CTH, as well as DMDS and SELE (**Fig. S9C**).

##### LncRNA interaction analysis

The known interactions of the enhancer lncRNAs in oligodendrocytes revealed that DLEU1 demonstrates significant interactions with a variety of biomolecules. Notably, it associates with DNA at the 13q14.3 locus and interacts with key proteins such as Argonaute 2 (AGO2) and mammalian target of rapamycin (MTOR). Furthermore, DLEU1 engages with the microRNAs hsa-miR-1271-3p and hsa-miR-200b-5p. It also forms complexes with RBPs IGF2BP2 and UPF1, as well as TFs FUS and AEBP2, highlighting its multifaceted role in cellular processes (**Fig. S9D, Table S13**).

#### ii. Astrocytes

The highest number of positively correlated lncRNA-mRNA pairs was observed in astrocytes. The chromosomal distribution of the lncRNAs reveals that Chr2 harbors the most, with 52 lncRNAs, followed by Chr1 (49), and Chr3 (41) (**Fig. S10A**). The lncRNA TTTY2B from the Y chromosome was positively correlated with 243 different mRNAs. Additionally, the lncRNAs AL353751.1, AL513128.2, AP000977.1, GALNT6, AC125616.1, AC012236.1, AC090282.1, AC243654.3, BABAM1, AC007405.1, AP001062.2, AC068631.1, LOC220729, AC068631.1, AC116351.1, AC116351.2, LINC02570, XLOC_006320, AC091729.3, AC011586.2, G080188, and SLA were found in close proximity to their correlated mRNA pairs (**Fig. S10B**).

##### Functional analysis of lncRNAs

The functional analysis of the enhancer lncRNAs demonstrated a significant enrichment in several biological pathways and activities, including RNA degradation pathways, protein O-linked glycosylation associated with threonine residues, methyltransferase activity, and RNA exonuclease activity, as well as cellular components related to both late and early endosomes (**Fig. S10C-F**).

##### Functional analysis of mRNAs

The mRNAs that are positively associated with the lncRNAs showed an enrichment of pentose phosphate pathway, RNA degradation, and SNARE interactions in vesicular transport pathways, biological processes including ribosome biogenesis, rRNA metabolic process, neuronal ion channel clustering, and molecular functions such as ion channel regulator activity and intramolecular oxidoreductase activity. There was also notable enrichment in the cellular complexes associated with the glutamatergic synapse and the peptidase complex (**Fig. S10G-J**).

##### Tissue-wise expression analysis

The tissue-level expression analysis of the positively regulated mRNAs indicates that most of them are predominantly expressed in the brain, including VGF, SLC6A17, and UQCRH (**Fig. S11A**). The analysis of lncRNA expression revealed that MEG3, BABAM1, INPP1, and COMMD1 demonstrated high expression levels across all examined brain regions. In contrast, LINC00470, MTUS2-AS2, and LINC00588 exhibited significantly lower expression levels (**Fig. S11B**).

##### PPI network analysis

The network of hub genes identified among the upregulated mRNAs depicts the protein-protein interactions involving TRIP13, HORMAD1, and MND1. Additionally, it highlights the interaction network among OTX2-FGF13-IFNA2-CD274 and UBL4A-GPI-EFL1-DDX51 (**Fig. S11C**).

##### PsyGeNET analysis

We have identified several of the upregulated genes to be linked with various psychiatric disorders. A total of 26 genes are positively associated with depressive disorders, 18 genes linked to bipolar disorder, and 14 genes correlated with alcohol use disorder (**Fig. S11D**).

##### LncRNA interaction analysis

The interaction analysis of the enhancer lncRNA in astrocytes demonstrates that DLX6-AS1, HAS2-AS1, AC135048.3, and LINC00343 engage in interactions with a variety of biomolecules. Notably, the CTCF TF interacts with multiple lncRNAs, including AC010503.1, AC113383.1, AC125616.1, AC135048.3, DLX6-AS1, HAS2-AS1, LINC00484, LINC00909, LINC01091, and VCAN-AS1. Additionally, the transcription factor FOXA1 is observed to interact with AC004895.1, LINC00343, LINC01091, LINC02217, and VCAN-AS1. The protein POLR2A is shown to engage with 12 distinct lncRNAs, specifically AC004895.1, AC010503.1, AC113383.1, AC125616.1, AC135048.3, DLX6-AS1, HAS2-AS1, LINC00484, LINC00909, LINC01091, LINC02217, and VCAN-AS1 (**Fig. S11E, Table S14**).

## Supplementary Figures

**Fig. S1:**
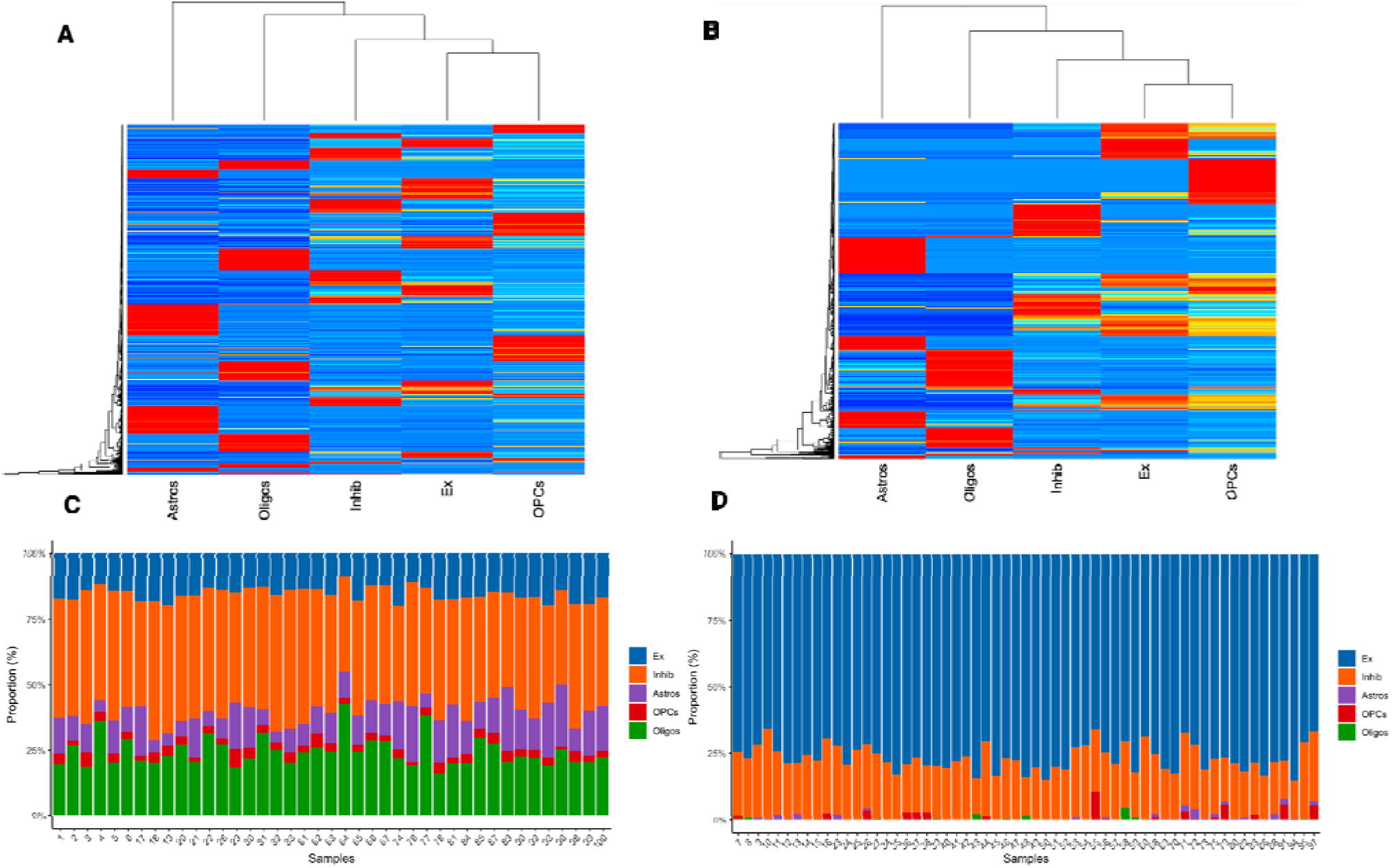
CIBERSORTx signature matrix and cell type proportions in Control and MDD subjects. **A-B** CIBERSORTx signature matrix heatmaps for A Control and B MDD conditions are displayed. Each row represents an individual gene, while each column corresponds to a major brain cell type, including astrocytes (Astros), oligodendrocytes (Oligos), inhibitory neurons (Inhib), excitatory neurons (Ex), and oligodendrocyte precursor cells (OPCs). Gene expression values are Z-score normalized. Hierarchical clustering effectively highlights gene expression patterns unique to each cell type. **C-D** Stacked bar plots of cell type proportions inferred by CIBERSORTx for C Control and D MDD samples. Each bar represents an individual sample, with colors indicating different brain cell types. The relative abundance of cell types varies between samples, with differences between control and MDD groups.

**Fig. S2:**
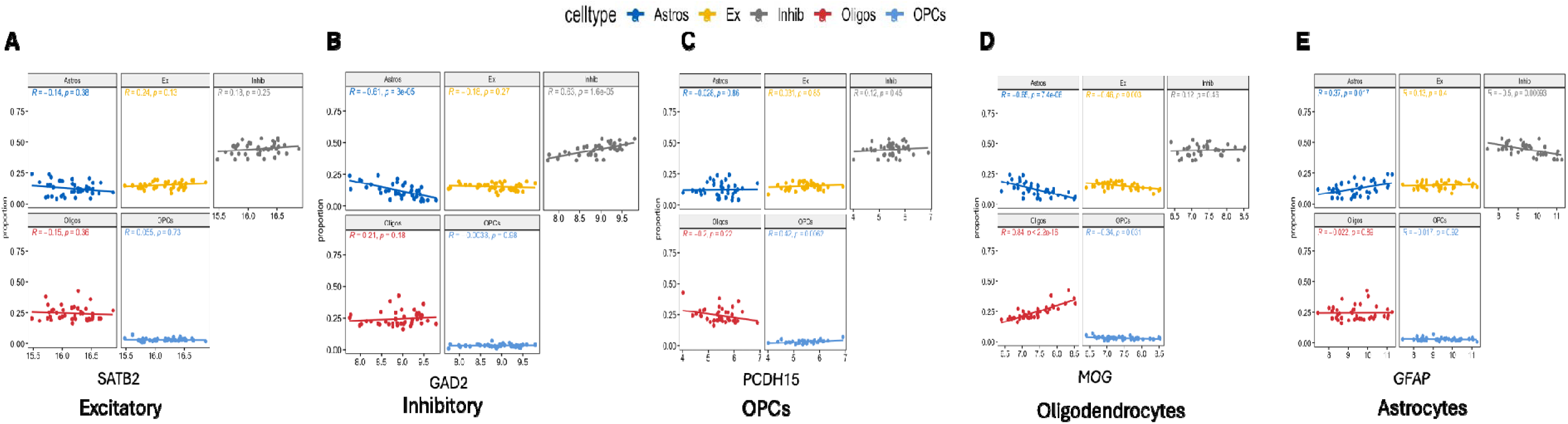
The Spearman correlation between the predicted cell type proportions and the expression levels of five marker genes (*SATB2, GAD2, PCDH15, MOG*, and *GFAP*) across five different brain cell types: Excitatory neurons, Inhibitory neurons, Oligodendrocyte Precursor Cells (OPCs), Oligodendrocytes, and Astrocytes.

**Fig. S3:**
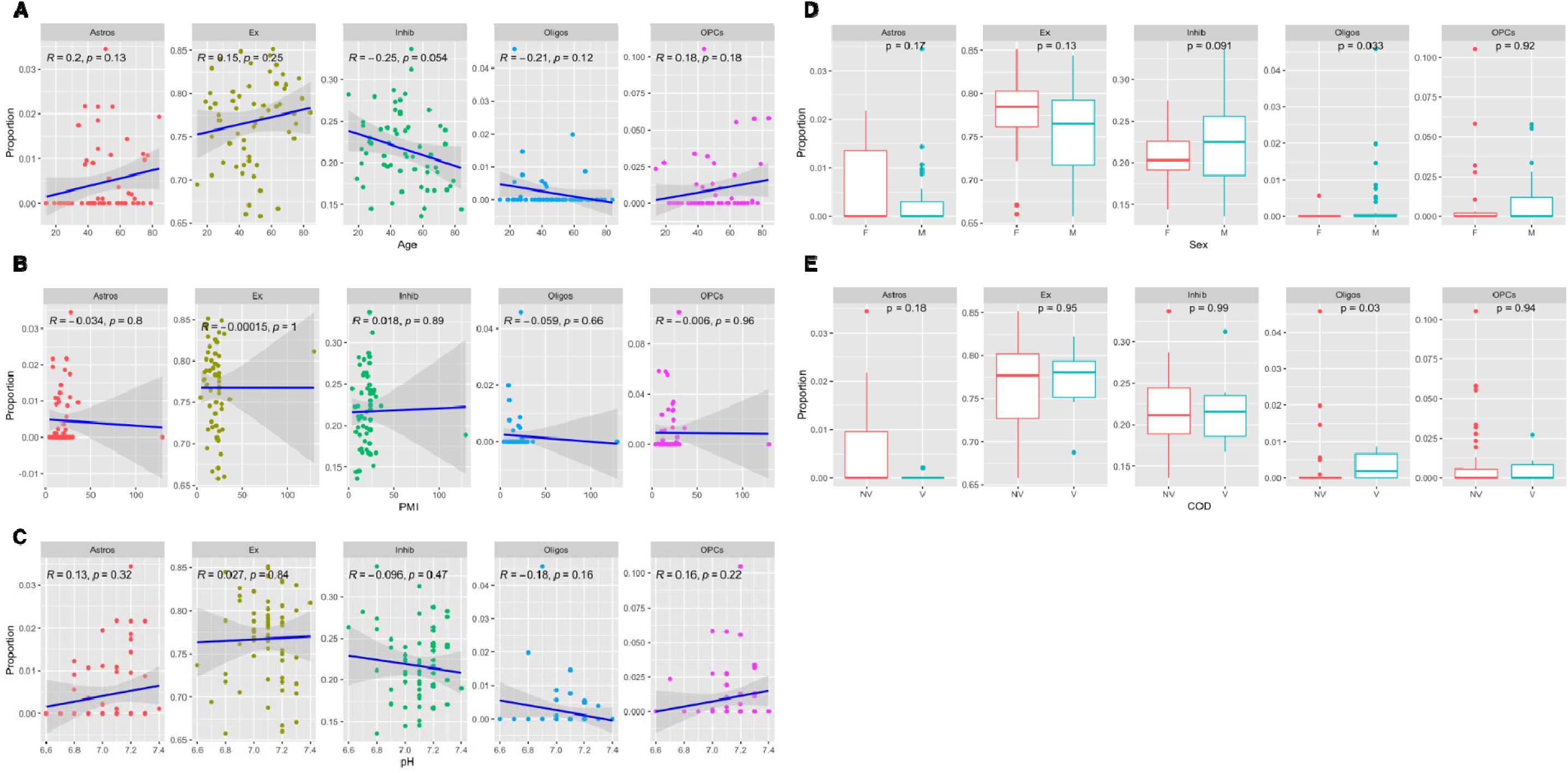
The association between predicted cell type proportions and various demographic factors, including **A** age, **B** post-mortem interval (PMI), **C** pH, **D** sex, and **E** cause of death (COD). Age and sex differences were significantly associated with the Oligodendrocytes proportion in the MDD. All other demographic features showed no significant association with proportions of other cell types.

**Fig. S4:**
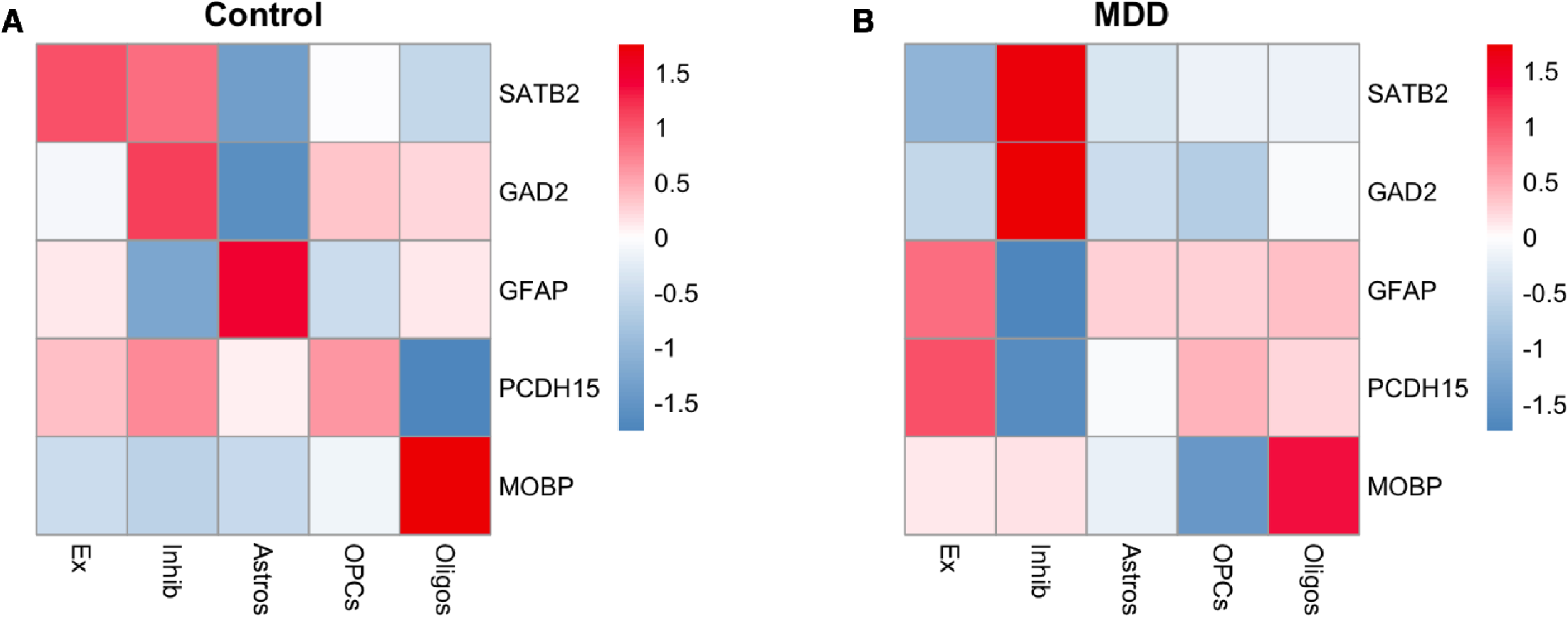
Correlation analysis of cell-type marker gene expression derived from single-nuclei RNA sequencing and deconvoluted profiles generated by bMIND. The genes *SATB2* (excitatory neurons), *GAD2* (inhibitory neurons), *GFAP* (astrocytes), *PCDH15* (OPCs), and *MOBP* (oligodendrocytes) exhibited the strongest correlations, indicating robust association between the two data sources.

**Fig. S5:**
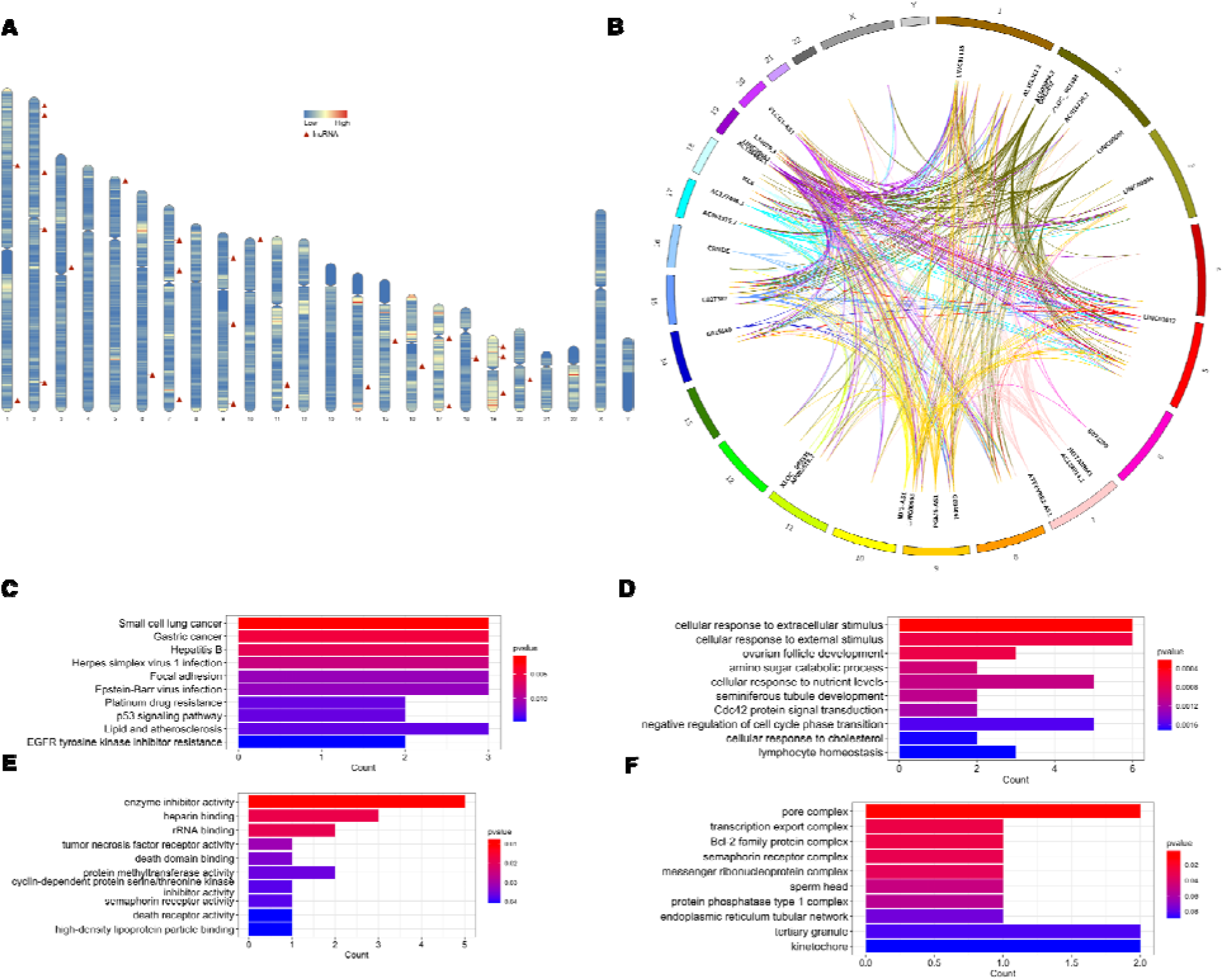
The inhibitory activity of lncRNAs on mRNAs, and the regulatory effects of lncRNAs in excitatory neurons. **A** Chromosomal distribution of the upregulated lncRNAs with negative regulatory effect on mRNAs identified in excitatory neurons. The karyotype is depicted with blue and red indicating low and high gene density, respectively. **B** The circos plot depicting the genomic distribution of negatively correlated upregulated lncRNAs and downregulated mRNAs in excitatory neurons. The outer ring represents the chromosomes, while the inner connections indicate pairs of correlated lncRNAs and mRNAs. **C** KEGG pathway enrichment of the mRNAs. **D** Biological Process GO term enrichment of the mRNAs. **E** Molecular Function GO term enrichment of the mRNAs. **F** Cellular Component GO term enrichment of the mRNAs.

**Fig. S6:**
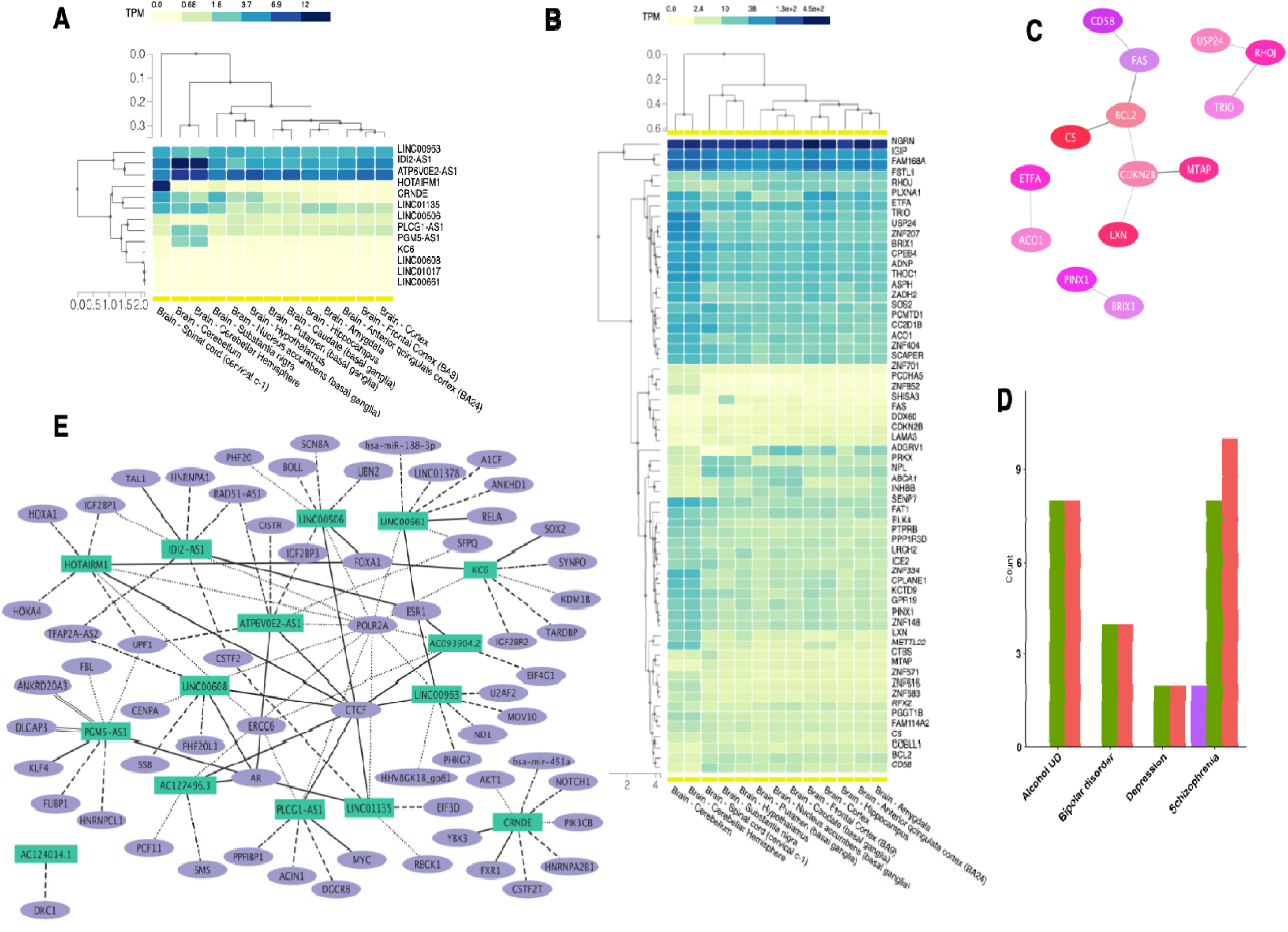
Functional characterization of the dysregulated correlating pairs of lncRNA and mRNAs in excitatory neurons exhibiting inhibitory activity. **A** A heatmap delineating the expression levels (TPM) of selected lncRNAs originating from excitatory neurons across various brain areas, as obtained from the GTEx database. **B** A heatmap of the selected mRNAs originating from excitatory neurons across various brain tissues. **C** Major protein-protein interaction networks of the negatively regulated mRNAs in excitatory neurons. **D** The number of genes associated with various neurological disorders from PsyGeNET database, including Alcohol Use Disorder, Bipolar Disorder, Cocaine Use Disorder, Depression, and Schizophrenia, identified among the negatively regulated mRNAs in excitatory neurons. 100% association shows that the genes are positively linked to the disorder. Both means there are reports of both positive and no association. **E** Network representation of known interactions between the top lncRNAs in inhibitory neurons and DNA, RNA, and proteins from the RNAInter database. Only the top two interactions with the highest score from each category are included. In the diagram, interactions with DNA are represented as parallel lines, RNA interactions as dash-dot lines, protein interactions as dots, RNA binding proteins as dashed lines, and transcription factors as solid lines.

**Fig. S7:**
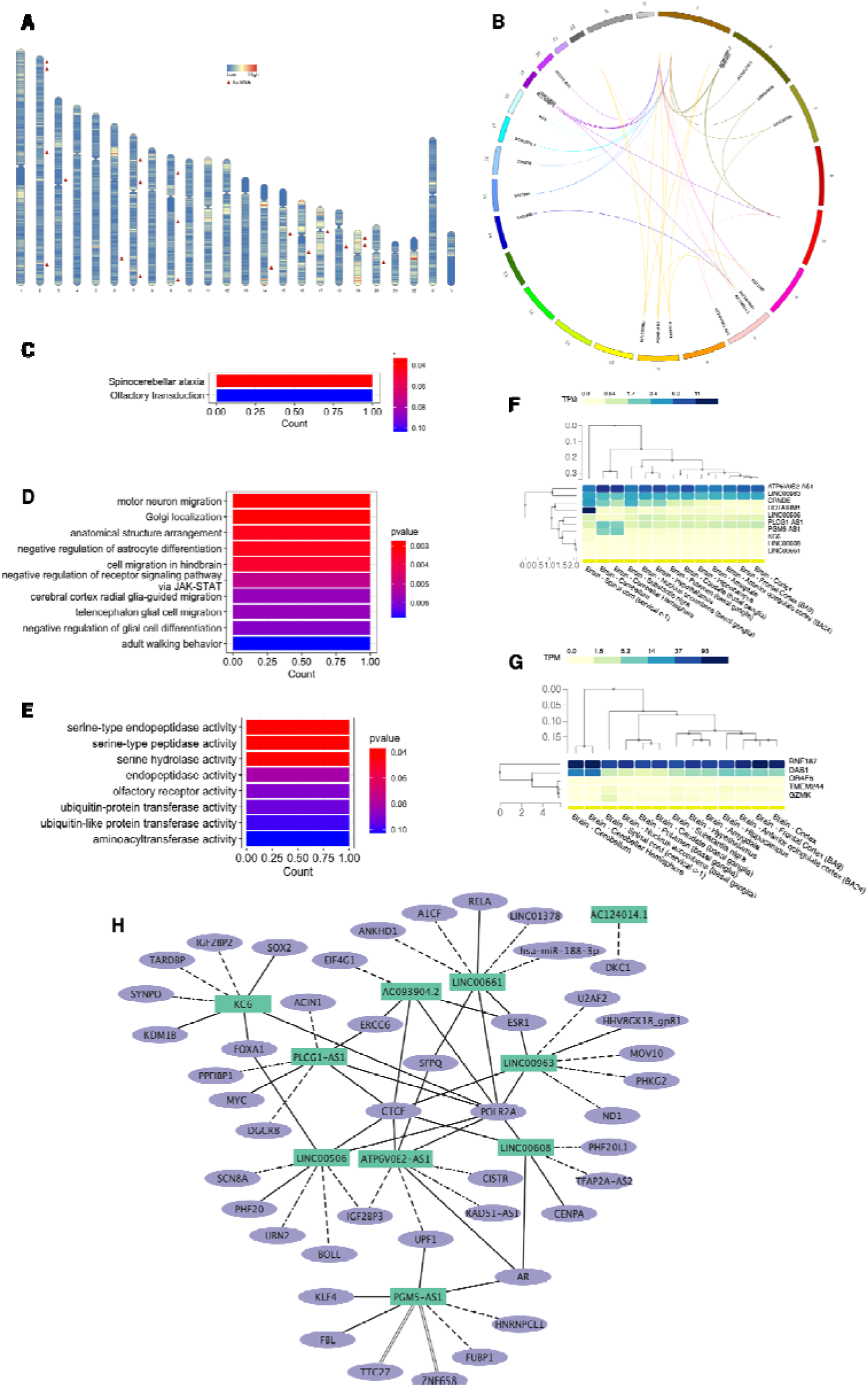
The analysis of the enhancer activity of lncRNAs on mRNAs and functional characterization in excitatory neurons. **A** Chromosomal distribution of the upregulated lncRNAs with positive regulatory effect on mRNAs identified in excitatory neurons. The karyotype is depicted with blue and red indicating low and high gene density, respectively. **B** The circos plot depicting the genomic distribution of positively correlated, upregulated lncRNAs and upregulated mRNAs in excitatory neurons. The outer ring represents the chromosomes, while the inner connections indicate pairs of correlated lncRNAs and mRNAs. The functional analysis of the positively correlated lncRNAs in excitatory neurons. **C** KEGG pathway enrichment of the mRNAs. **D** Biological Process GO term enrichment of the mRNAs. (**E)** Molecular Function GO term enrichment of the mRNAs. **F** A heatmap delineating the expression levels (TPM) of selected lncRNAs originating from excitatory neurons across various brain regions, as obtained from the GTEx database. **G** A heatmap of the selected mRNAs originating from excitatory neurons across various brain tissues. **H** Network representation of known interactions between the top lncRNAs from excitatory neurons and DNA, RNA, and proteins from the RNAInter database. Only the top two interactions with the highest score from each category are included. In the diagram, interactions with DNA are represented as parallel lines, RNA interactions as dash-dot lines, protein interactions as dots, RNA binding proteins as dashed lines, and transcription factors as solid lines.

**Fig. S8:**
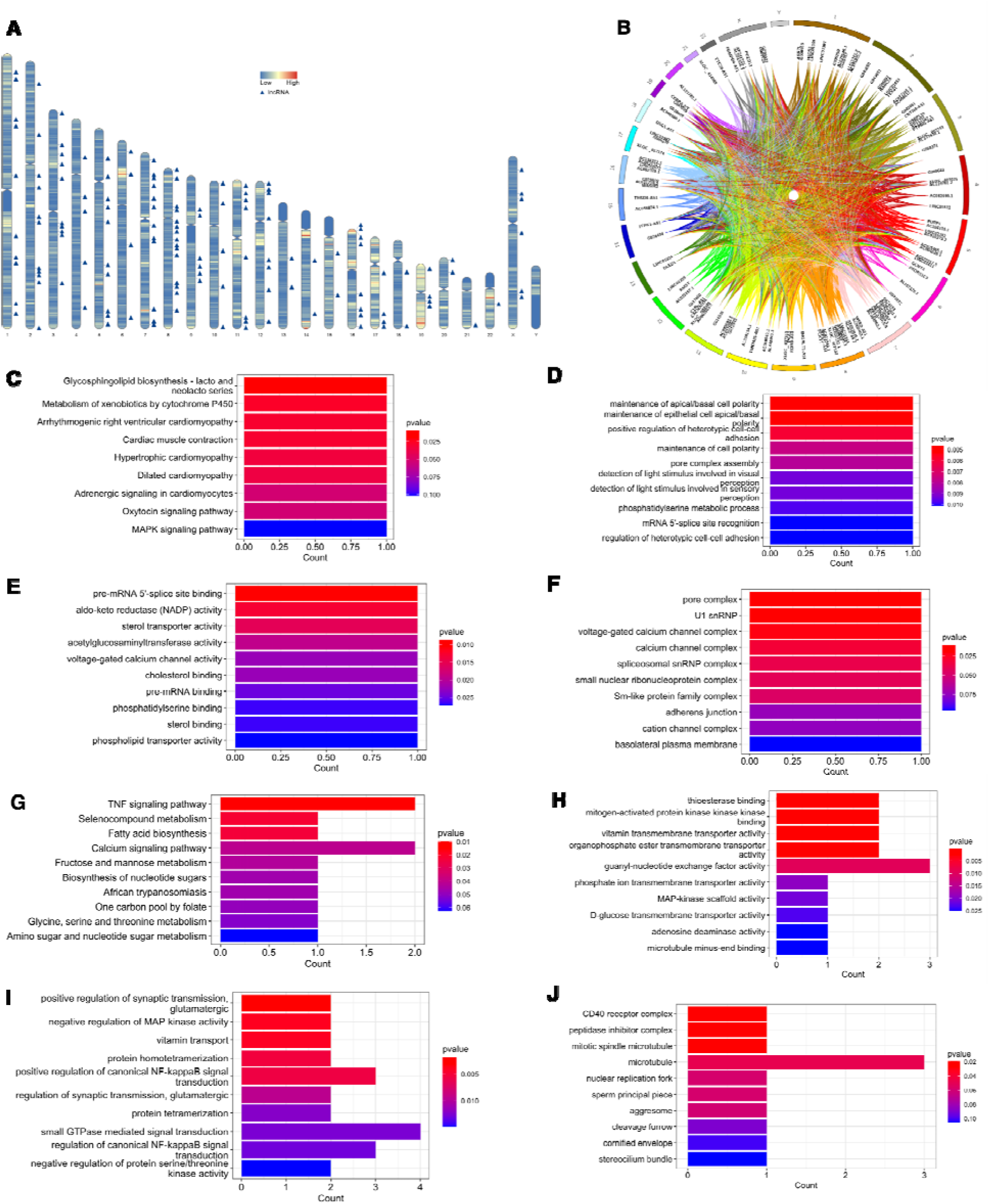
The analysis of the enhancer activity of lncRNAs on mRNAs and functional characterization in oligodendrocytes. **A** Chromosomal distribution of the upregulated lncRNAs with positive regulatory effect on mRNAs identified in oligodendrocytes. The karyotype is depicted with blue and red indicating low and high gene density, respectively. **B** The circos plot depicting the genomic distribution of positively correlated, upregulated lncRNAs and upregulated mRNAs in oligodendrocytes. The outer ring represents the chromosomes, while the inner connections indicate pairs of correlated lncRNAs and mRNAs. The functional analysis of the positively correlated lncRNAs and mRNAs in oligodendrocytes. **C** KEGG pathway enrichment of the lncRNAs. **D** Biological Process GO term enrichment of the lncRNAs. **E** Molecular Function GO term enrichment of the lncRNAs. **F** Cellular Component GO term enrichment of the lncRNAs. **G** KEGG pathway enrichment of the mRNAs. **H** Biological Process GO term enrichment of the mRNAs. **I** Molecular Function GO term enrichment of the mRNAs. **J** Cellular Component GO term enrichment of the mRNAs.

**Fig. S9:**
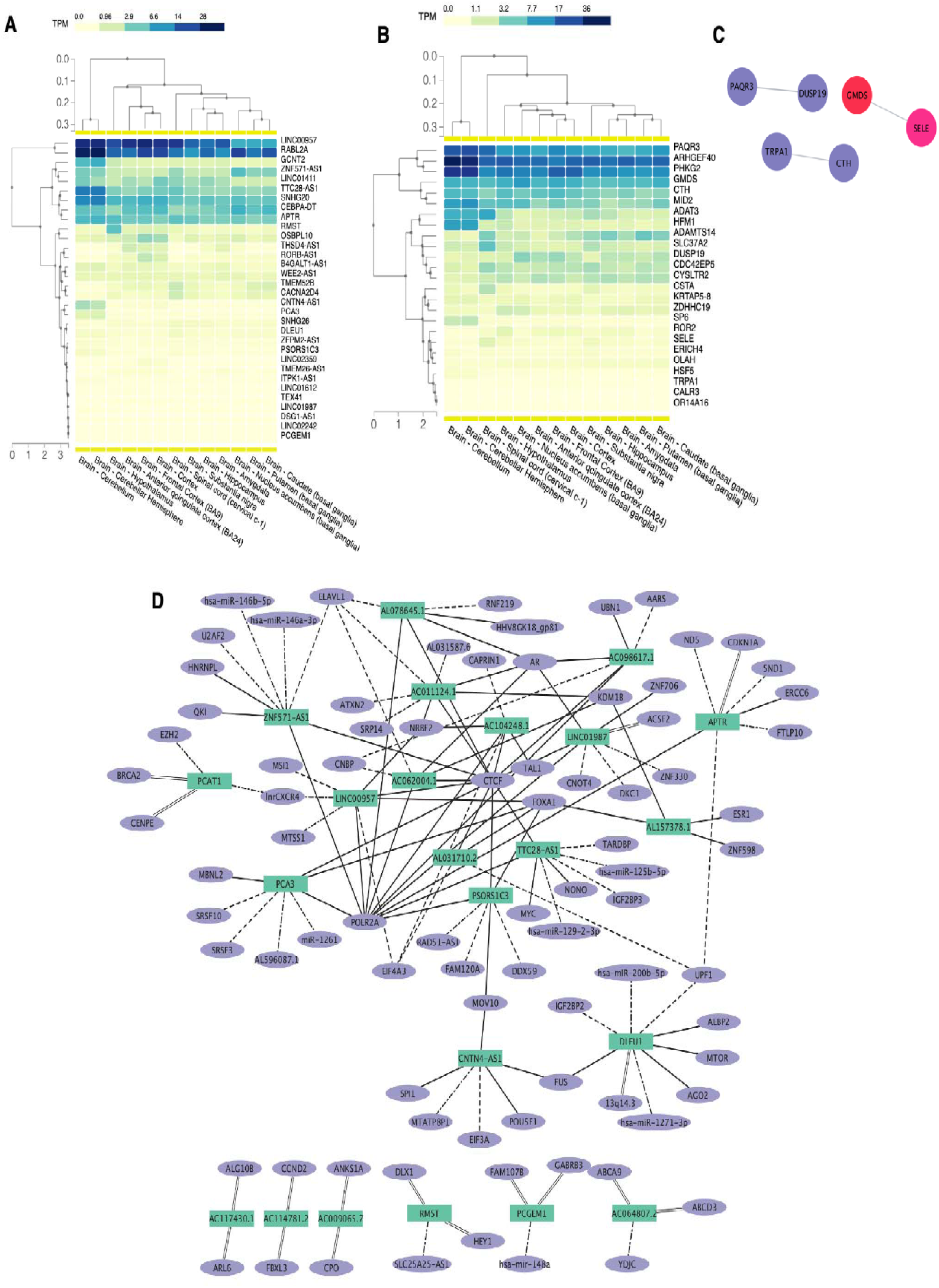
Functional characterization of the dysregulated correlating pairs of lncRNA and mRNAs in oligodendrocytes exhibiting enhancer activity. **A** A heatmap depicting the expression levels (TPM) of selected lncRNAs originating from oligodendrocytes across various brain regions, as obtained from the GTEx database. **B** A heatmap of the selected mRNAs originating from oligodendrocytes across various brain tissues. **C** Major protein-protein interaction networks of the positively regulated mRNAs in oligodendrocytes. **D** Network representation of known interactions between the top lncRNAs from oligodendrocytes and DNA, RNA, and proteins from the RNAInter database. Only the top two interactions with the highest score from each category are included. In the diagram, interactions with DNA are represented as parallel lines, RNA interactions as dash-dot lines, protein interactions as dots, RNA binding proteins as dashed lines, and transcription factors as solid lines.

**Fig. S10:**
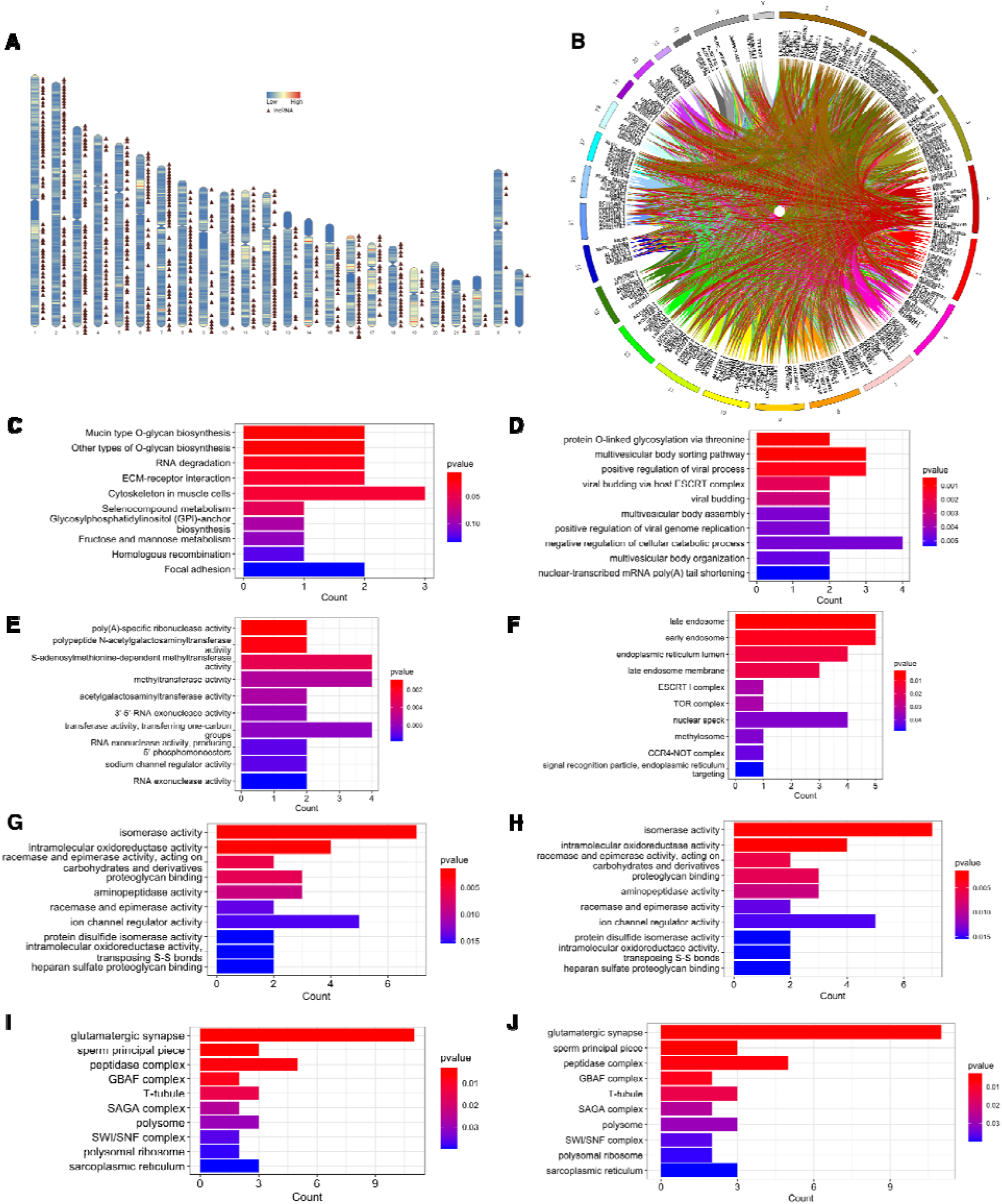
The analysis of the enhancer activity of lncRNAs on mRNAs and functional characterization in astrocytes. **A** Chromosomal distribution of the upregulated lncRNAs with positive regulatory effect on mRNAs identified in astrocytes. The karyotype is depicted with blue and red indicating low and high gene density, respectively. **B** The circos plot depicting the genomic distribution of positively correlated upregulated lncRNAs and upregulated mRNAs in astrocytes. The outer ring represents the chromosomes, while the inner connections indicate pairs of correlated lncRNAs and mRNAs. The functional analysis of the positively correlated lncRNAs and mRNAs in astrocytes. **C** KEGG pathway enrichment of the lncRNAs. **D** Biological Process GO term enrichment of the lncRNAs. **E** Molecular Function GO term enrichment of the lncRNAs. **F** Cellular Component GO term enrichment of the lncRNAs. **G** KEGG pathway enrichment of the mRNAs. **H** Biological Process GO term enrichment of the mRNAs. **I** Molecular Function GO term enrichment of the mRNAs. **J** Cellular Component GO term enrichment of the mRNAs.

**Fig. S11:**
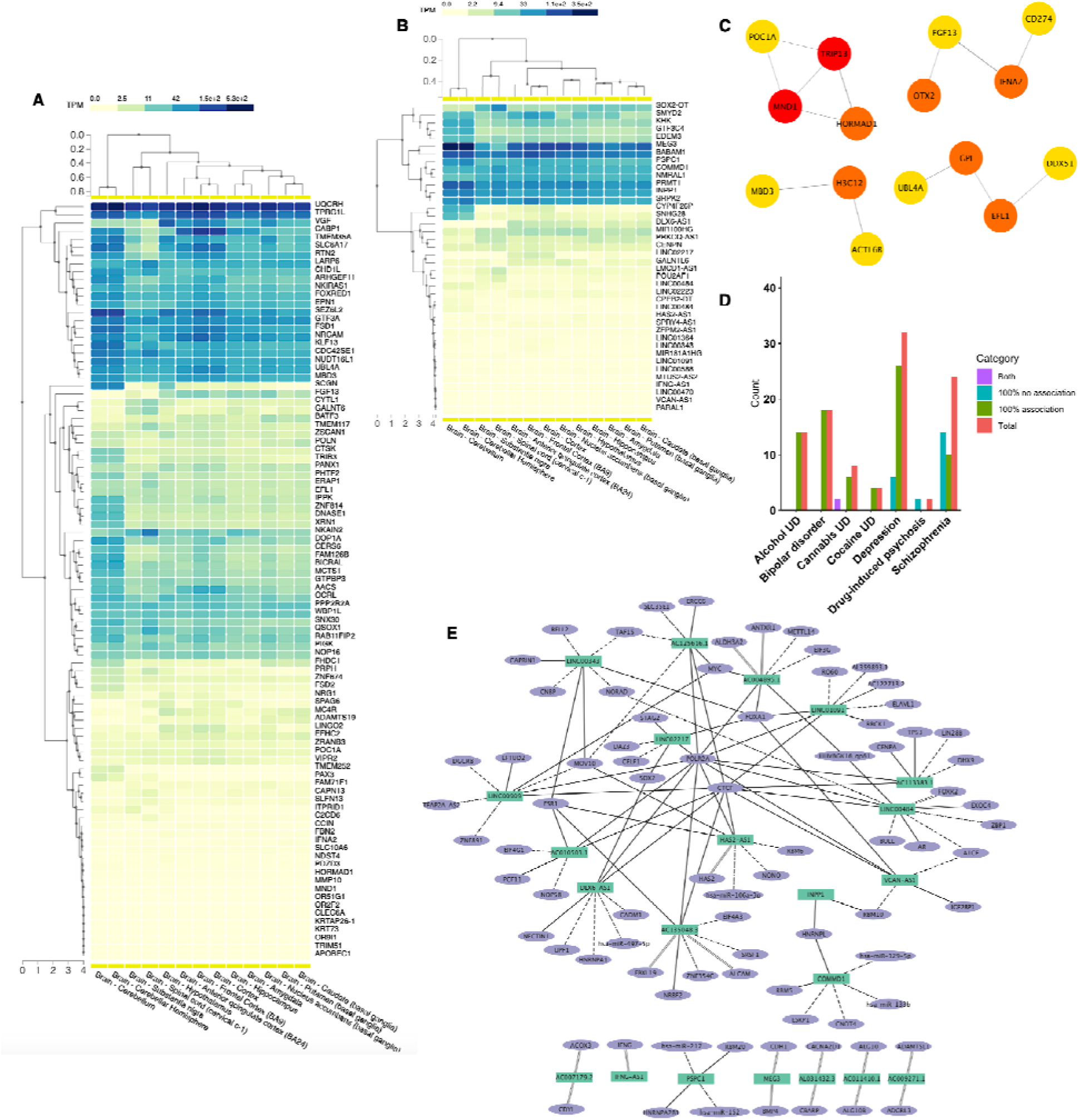
Functional characterization of the dysregulated correlating pairs of lncRNA and mRNAs in astrocytes exhibiting enhancer activity. **A** A heatmap delineating the expression levels (TPM) of selected mRNAs originating from astrocytes across various brain regions, as obtained from the GTEx database. **B** A heatmap of the selected lncRNAs originating from astrocytes across various brain tissues. **C** Major protein-protein interaction networks of the hub genes from positively regulated mRNAs in astrocytes. **D** The number of genes associated with various neurological disorders from PsyGeNET database, including Alcohol Use Disorder, Bipolar Disorder, Depression, and Schizophrenia, identified among the positively regulated mRNAs in inhibitory neurons. 100% no association indicates that the genes are not linked to the disorder, while 100% association shows that the genes are positively linked to the disorder. Both means there are reports of both positive and no association. **E** Network representation of known interactions between the top lncRNAs from astrocytes and DNA, RNA, and proteins from the RNAInter database. Only the top two interactions with the highest score from each category are included. In the diagram, interactions with DNA are represented as parallel lines, RNA interactions as dash-dot lines, protein interactions as dots, RNA binding proteins as dashed lines, and transcription factors as solid lines.

## Supplementary Table Legends

**Table S1:** Demographic and clinical characteristics of subjects.

| **Table S1: Demographic and clinical characteristics of subjects** |  |  |
| --- | --- | --- |
|  | **Non-psychiatric Controls** | **MDD Subjects** |
| **Number of subjects** | 41 | 59 |
| **Age (Year)** | 49.46±2.84 | 47.71±2.26<br>(F=0.09, p=0.63, t=0.48, df=98) |
| **PMI (Hours)** | 18.26±0.92 | 20.66±2.07<br>(F=1.54, p=0.36, t=-0.92, df=98) |
| **RIN** | 7.68±0.04 | 7.07±0.02<br>(F=0.44, p=0.54, t=0.61, df=98) |
| **Brain pH** | 7.10±0.03 | 7.27±0.23<br>(F=2.63, p=0.40, t=0.84, df=98) |
| **Gender** |  |  |
| **Males** | 26 | 35 |
| **Females** | 15 | 24 |
| **Cause of Death** | Pneumonia, accidental chest injury, atherosclerotic cardiovascular disease, cardiac arrhythmia, multiple vehicle accident, embolism, leukemia, morbid obesity, electrocution, multiple injuries, GSW, respiratory failure, cardiopulmonary arrest, complications due to diabetes, upper GI bleed, cardiopulmonary arrest, colon cancer, renal failure | GSW, jumped from height, hanging, CO intoxication, drug overdose, stab wound, atherosclerotic cardiovascular disease, multiple vehicle accident, ketoacidosis, cardiomegaly, seizure, fatty liver, cardiac arrhythmia, hemopericardium, liver failure, leukemia, pulmonary embolism, lymphoma, lung cancer, acute myocardial infarction |
| **Race** | 5 Black/36 White | 6 Black/1 Asian/52 White |
| **Neurological/**<br>**Neuropathological disorders** | None | None |
| **Number of subjects showing**<br>**positive antidepressant toxicology** | None | 31 |
| **Number of subjects showing**<br>**substance use** | None | 4 |
| Data are the mean ± SEM; The data were analyzed using Independent Sample t-Test. MDD group was compared with the control group. |  |  |

**Table S2: lncRNA microarray expression data.** Column A: ProbeName, it represents the probe name, Column B ∼ CW: Raw Intensity of each sample, Column CX ∼ GS: Normalized Intensity of each sample (transformed), Column GT ∼ KO: Flags, flags are attributes that denote the quality of the entities (auto-flagging by GeneSpring GX v12.1 software), Column KP ∼ LY: Annotations to each probe, including Transcript_ID, GeneID, GeneSymbol, type, LncRNA_Completeness, DHS_type, geneCategory, lncRNA_level, Validated, Canonical, Locus, chrom, strand, txStart, txEnd, CancerName, DiseaseName, associated_cell_or_tissue, Subcellular_Localization, source, description, RNAlength, Sequence, TIR_conservation, exon_conservation, TraitName, eQTL_cis_correlated_mRNA_geneID, smProtein, refseqID, relationship, Associated_gene_acc, Associated_gene_name, Associated_protein_name, Associated_gene_strand, associated_gene_start and Associated_gene_end.

**Table S3: mRNA microarray expression data.** Column A: ProbeName, it represents the probe name, Column B ∼ CW: Raw Intensity of each sample, Column CX ∼ GS: Normalized Intensity of each sample (transformed), Column GT ∼ KO: Flags, flags are attributes that denote the quality of the entities (auto-flagging by GeneSpring GX v12.1 software), Column KP ∼ LN: Annotations to each probe, including Transcript_ID, GeneID, GeneSymbol, type, LncRNA_Completeness, DHS_type, geneCategory, lncRNA_level, Validated, Canonical, Locus, chrom, strand, txStart, txEnd, CancerName, DiseaseName, associated_cell_or_tissue, Subcellular_Localization, source, description, RNAlength, Sequence, TIR_conservation, exon_conservation, TraitName, eQTL_cis_correlated_mRNA_geneID, smProtein, refseqID, relationship, Associated_gene_acc, Associated_gene_name, Associated_protein_name, Associated_gene_strand, associated_gene_start and Associated_gene_end.

**Table S4: lncRNA Differential expression.** Columns A∼F: Excitatory neurons, H∼M: Inhibitory neurons, O∼T: OPCs, V∼AA: Oligodendrocytes, AC∼AH: Astrocytes.

**Table S5: mRNA differential expression.** Columns A∼F: Excitatory neurons, H∼M: Inhibitory neurons, O∼T: OPCs, V∼AA: Oligodendrocytes, AC∼AH: Astrocytes.

**Table S6: Inhibitory regulation of lncRNAs on mRNAs.** Columns A∼D: Excitatory neurons, F∼I: Inhibitory neurons.

**Table S7: Cellular localization of Inhibitory lncRNAs.** Columns A∼H: Inhibitory neurons, J∼Q: Excitatory neurons.

**Table S8:** The known interactions of the Inhibitory lncRNAs in Inhibitory neurons. Columns A∼M: DNA interactions, O∼AA: RNA interactions, AC∼AO: Protein interactions.

**Table S9:** The known interactions of the Inhibitory lncRNAs in Excitatory neurons. Columns A∼M: DNA interactions, O∼AA: RNA interactions, AC∼AO: Protein interactions.

**Table S10: Positive regulation of lncRNAs on mRNAs in different cell types.** Columns A∼D: Excitatory neurons, F∼I: Inhibitory neurons. K∼N: Oligodendrocytes, P∼S: Astrocytes.

**Table S11:** The known interactions of the enhancer lncRNAs in Inhibitory neurons. Columns A∼M: DNA interactions, O∼AA: RNA interactions, AC∼AO: Protein interactions.

**Table S12:** The known interactions of the dysregulated lncRNAs in Excitatory neurons. Columns A∼M: DNA interactions, O∼AA: RNA interactions, AC∼AO: Protein interactions.

**Table S13: The known interactions of the enhancer lncRNAs in Oligodendrocytes.** Columns A∼M: DNA interactions, O∼AA: RNA interactions, AC∼AO: Protein interactions.

**Table S14: The known interactions of the enhancer lncRNAs in Astrocytes.** Columns A∼M: DNA interactions, O∼AA: RNA interactions, AC∼AO: Protein interactions.

